## Supplementary material for "The new archaeal order Lutiacidiplasmatales reveals convergent evolution in Thermoplasmatota": SI text

### Supplementary Methods

#### *Extended Phylogenomics*

##### *Datasets*

This study used three datasets to build a full Thermoplasmatota tree, a Thermoplasmatota tree containing only high-quality genomes, and a tree restricted to the Lutiacidiplasmatales.

The **full dataset** tree comprises 124 archaeal genomes with completeness greater than 45% and less than 10% contamination. This dataset contains the 35 newly sequenced genomes, the TMEG-bg1 genome, 84 genomes representing 84 other Thermoplasmatota species and 4 Archaeoglobales genomes (the outgroup).

The **high-quality dataset** tree comprises 100 archaeal genomes with completeness greater than 70% and less than 5% contamination. This dataset contains 21 newly sequenced genomes, the TMEG-bg1 genome, 74 genomes representing 74 other Thermoplasmatota species and four Archaeoglobales genomes.

The **Lutiacidiplasmatales-specific dataset** comprises 40 archaeal genomes with completeness greater than 45% and less than 10% contamination. This dataset contains 35 newly sequenced genomes, two publically available Lutiacidiplasmatales genomes and 3 Methanomassiliicoccales genomes (the outgroup).

##### *Ortholog selection*

For each dataset, ortholog groups (OGs) were detected using Roary (-i 50, -iv 1.5)[1]. Core OGs were defined as those present in a single copy in each genome and present in at least 50% of the genomes for the full dataset and the Lutiacidiplasmatales-specific dataset, or 70% of genomes in the high-quality dataset. Core OGs were aligned individually using MAFFT L-INS-i [2], and spurious sequences and poorly aligned regions were removed with trimAl (automated 1, resoverlap 0.55 and seqoverlap 60)[3]. Alignments were removed from further analysis if they presented evidence of recombination using the PHItest[4]. The remaining alignments from each of the three datasets were concatenated into supermatrices.

*Estimating phylogeny:* To establish a robust phylogeny of the Thermoplasmatota, we compared seven phylogenomic trees reconstructed using different approaches. The first four trees were estimated by selecting gene markers from two taxonomic samplings (the full and high-quality datasets) (Figure S24). Maximum likelihood trees were constructed for each supermatrix of alignments with IQ-TREE 2.0.3[5], using the best fitting model in ModelFinder[6] for each alignment and an edge-linked partition model. The same two alignments were also subjected to an additional round of trimal (automated1) on the supermatrix before using the mixture model LG+C60+F model, resulting in four species trees. Branch validation of each tree involved 1000 SH-aLRT test[7] and 2000 UFBoot replicates, and a hill-climbing nearest neighbour interchange (NNI) search was performed to reduce the risk of overestimating branch supports. A fifth tree was constructed using a concatenation of 17 ribosomal genes from the full dataset using the best fitting model in ModelFinder[6] for each alignment and an edge-linked partition model.

Single-gene trees were constructed for the 71 marker gene alignments from the high-quality genome dataset using IQ-TREE 2.0.3 [5] and the best fitting model in ModelFinder[6]. These single-gene trees were used to construct a supertree using the multispecies coalescence method implemented in ASTRAL v5.7.5[8]. A phylogenomic tree was also constructed for the full dataset concatenated supermatrix after SR4 recoding[9], using IQ-TREE 2.0.3[5] and the best fitting model in ModelFinder[6].

*Topology testing in constraint trees:* ML trees were created for the high-quality dataset to form trees that with the topology unconstrained, constrained to place Posedoniales and Thermoprofundales as basal paraphyletic groups of the Thermoplasmatota (as in Adam et al 2017[10]) or as a basal monophyletic group (as was suggested, albeit with poor support, by the SR4 recoded tree). These topologies were then compared with approximately unbiased test[11] and other statistical tests implemented in IQ-TREE 2.0.3 [5] (-zb 10000 -zw -au) (Figure S4).

***Functional annotation of gene families***

Carbohydrate active enzymes were annotated using profile HMM from dbCAN (http://bcb.unl.edu/dbCAN2/) (filtered with hmmscan-parser.sh and by removing matches with mean posterior probability <0.7). Extracellular peptidases were initially annotated using Pfam profile HMMs corresponding to MEROPs families, as described by Tully et al. [12], to identify peptidases and then predict signal peptides' presence in these genes using Signalp 5.0[13] (-org arch, archaeal signal peptides).

The presence of motility genes in Thermoplasmatota was initially assessed by the presence of the conserved archaellum subunits C (arCOG05119), D/E (arCOG02964), F (arCOG01824), G (arCOG01822) and J (arCOG01809). However, Tully et al. indicated that Poseidoniales species might possess divergent motility loci[12]. Therefore, profile HMMs of archaeal flagellin (PF01917) and *flaH* (PF06745) genes were used as markers for possible divergent motility, even when other archaellum related genes were absent.

The subfamily classification of *cyd*A was performed using hmmsearch with the *cyd*A subfamily database [14].

The subfamily classification of *coxA* genes was performed using the heme-copper oxygen reductase database [15].

***Sulfite oxidation genes***

Protein sequences possessing the Pfam oxidoreductase molybdopterin binding domain (PF00174) were downloaded from Swiss-Prot. They were combined with sequences containing the PF00174 conserved domain from the full Thermoplasmatota genome dataset (this study) and from the Thaumarchaeota genome dataset (Sheridan et al. 2020[16]), resulting in 213 sequences. Conserved domains within the sequences were annotated using InterProScan [17], Pfam [18], SUPERFAMILY [19] and MobiDBLite [20] databases. Transmembrane helices were predicted using TMHMM 2.0 [21].

The 213 sequences were then aligned using MAFFT L-INS-i [2], processed with trimAl (automated1)[3], and an ML phylogenetic tree was constructed using IQ-TREE 2.0.3[5] with 2,000 UFBoot replicates and 1,000 SH-aLRT test[7], an NNI search and the best substitution model selected by ModelFinder[6]). The ancestral deviation for each node was calculated using MAD programme[22], and the node with the minimal ancestor deviation was used as the tree's root.

***Note on gene family annotation***

A high novelty of the proteomes was observed. A high percentage of the gene families could not be functionally annotated at this time (Supplementary Data 18), with 50% of the gene families lacking close homologs in the arCOG database.

### Supplementary Results

***A robust phylogeny for the Thermoplasmatota. Short name: SI Phylogenomics***

The increased availability of genomes in recent years and the new genomes sequenced in this study allow revisiting deep evolutionary relationships within the Thermoplasmatota. Seven phylogenomic trees were created to estimate the phylogeny of the Thermoplasmatota using different approaches. All seven species trees constructed in this work for Thermoplasmatota differed to some extent from the topology presented in Adam *et al.* 2017[10] (a thorough investigation spanning multiple archaeal phyla) (Figure S2, Topology B) and some other works[23, 24]. Six trees (Trees 1-6 from Figure S1) resolved Acidiprofundales and Thermoplasmatales as a basal monophyletic group in the Thermoplasmatota (Figure S2, Topology A). In contrast, the remaining tree (Tree 6 from Figure S1) implied an internal branching of this group (albeit with very poor support) (Figure S3, Topology C). Constraint tree statistical analysis of these three differing topologies strongly favoured topology A and could statistically reject topology B. Trees with this topology were used in further evolutionary analysis. Marker gene information for the full dataset trees is provided in Supplementary Data 20.

***Distribution of posterior probabilities in predictions of gene family origination events***

The likelihood of each gene family originating a single time into the Thermoplasmatota was estimated for every candidate originating branch (Supplementary Data 21). With a 0.5 posterior probability (PP) criterion on a single branch, over 70% (4,256 of 6,050) of the gene families were predicted to have been acquired a single time into the Thermoplasmatales. This percentage declined to 60, 50, 38, 25 and 9 % when the threshold was increased to PPs greater than 0.6, 0.7, 0.8, 0.9 and equal to 1.0, respectively. Therefore, even at the permissive criterion (0.5 PP), a single point of origination could not be predicted for almost 30 % of the gene families used in the gene tree – species tree reconciliation analysis, and this number increased notably as the PP threshold was made more stringent (Figure S25).

***Origination and evolution of complex IV assembly components ctaA, ctaB and coxB. Short name: SI Complex IV evolution.***

The *cta*A genes detected in Lutiacidiplasmatales and Poseidonales were highly divergent from each other (Figure S11), indicating two independent acquisitions of *cta*A into the Thermoplasmatota. The *cta*A genes of Poseidonales were potentially acquired from cyanobacteria, given their *cta*A close phylogenetic relationship and their shared marine environment. In contrast, the Lutiacidiplasmatales *cta*A genes are affiliated with various bacterial lineages with no discernible shared environment.

The *cta*B gene, which is responsible for the biosynthesis of haem O from haem B, appears to have a more complicated evolutionary history. Again, the Poseidonales genes diverged from the other Thermoplasmatota and cluster more closely with bacterial homologs (Figure S12), indicating independent originations into Thermoplasmatota. In addition, the three orders, Thermoplasmatales, Lunaplasmatales and Lutiacidiplasmatales, were all separated into two clades. Gene tree - species tree reconciliation indicates that this ancestral splitting did not occur through an ancient duplication event. Therefore, it is likely that there were at least three independent origins of *cta*B in the Thermoplasmatota: one ancestral gene present in Thermoplasmatales, one gene which has possibly been acquired from the Thaumarchaeota and is present in Thermoplasmatales, Lutiacidiplasmatales and Lunaplasmatales, and a third of bacterial origin into the Poseidonales (Figure S12). The presence of *cta*B in SAL16 and TMEG-bg1 is suggested by gene tree - species tree reconciliation to have most likely (albeit with weak support) occurred by transfer from the Thermoplasmatales LCA to SAL16 (0.31 TPP (posterior probability of transfer)) and then subsequently from SAL16 to the LCA of TMEG-bg1 and UBA184 (0.26 TPP) (Supplementary Data 22).

In contrast to the other subunits of the complex IV, The *cox*B genes of Lutiacidiplasmatales, Poseidoniales, Lunaplasmatales and most Thermoplasmatales genomes cluster together. At the same time, a Thermoplasmatales clade, consisting of the genera *Ferroplasma*, *Acidiplasma* and *Picrophilus,* possess divergent *cox*B genes more similar to that of *Halobiforma lacisalsi* (Figure S12). The reason for this is unknown, but it is noteworthy that several of the organisms that possess this divergent *cox*B are acidophilic ferrous iron oxidisers from distantly related microbial lineages.

#### ***Progressive evolutionary history of Lutiacidiplasmatales***

The evolution of Lutiacidiplasmatales from the Thermoplasmatota LCA is predicted to have been marked by at least three bifurcating divergences. Functional gene gain and loss were analysed by comparing progressive ancestral genome reconstructions (Figure S26) and validated by origination posterior probability (OPP) if the gene family is predicted to have originated only once in Thermoplasmatota.

The first bifurcating divergence in this analysis formed a Thermoplasmatota clade, TP_2 LCA, excluding o_Acidiprofundales and o_Thermoplasmatales. This divergence coincided with the gain of gene families, including a nickel-containing superoxide dismutase, a K+ stimulated pyrophosphate-energised sodium pump and several amino acid metabolism genes, and the loss of gene families including the Pgi1-type glucose-6-phosphate isomerase.

The second divergence exclude the o_Poseidonales and o_Thermoprofundales. This divergence coincided with the gain of genes such as a divergent form of the oxidative protective peptide methionine sulfoxide reductase and phosphoglycolate phosphatase, which prevents the inhibition of glycolysis by phosphoglycolate. It also coincided with the loss of genes such as poorly characterised Archaeal PilT-family ATPase.

The third divergence comprises only the Lutiacidiplasmatales LCA, excluding the rest of the Thermoplasmatota lineages. This divergence coincided with one of the most significant gene family gains by origination detected in the Thermoplasmatota. Several genes involved in glycolysis were gained, including ATP-dependent phosphofructokinase *pfk*, and fructose-bisphosphate aldolase class I, *fba*B (0.97 OPP). Several gene families involved in oxidative phosphorylation were also acquired, including the originating gene families heme A synthase, *cta*A, and heme-copper oxygen reductase subunit b, *cox*B. Additionally, this divergence coincided with the origination of heterotrophy genes such as the sarcosine oxidase subunits A (0.87 OPP) and B (0.99 OPP) and the gain of pentose phosphate pathway genes glucose-6-phosphate 1-dehydrogenase, *zwf*, and 6-phosphogluconate dehydrogenase, gnd. The third divergence also included some notable losses, such as the three genes involved in the biosynthesis of histidine (*his*B, F and G) from products of the pentose phosphate pathway and the loss of indolepyruvate ferredoxin oxidoreductase (both A and B subunits lost), an enzyme involved in archaeal peptide fermentation[25]. The expanded functional gene gain and loss in the evolution from the Thermoplasmatota LCA to Lutiacidiplasmatales LCA can be found in Tables S23.

Gene families predicted to have originated in the LCA of Lutiacidiplasmatales were queried against the UniRef90 database using the permissive criteria (E-value $1x10^{-3}$). Homologs were detected for 81 % of the originating gene families. However, over half of these hits were classified with terms such as "Thermoplasmata archaeon" and "Euryarchaeota_archaeon" or "Mine drainage metagenome", which is an environment where Lutiacidiplasmatales is known to be prevalent, from the 16S rRNA gene analysis shown earlier. There was, therefore, a significant chance of some of the hits being for genes from Lutiacidiplasmatales or other Thermoplasmatota species. Therefore, UniRef90 was filtered for genes with strain-level classifications and genes from Thermoplasmatota species were removed. Homologs were detected for 55 % of gene families originating in the LCA of Lutiacidiplasmatales using the filtered database (Tables S18). Fifty-one per cent of these hits were for genes from bacteria, and 49 % were for genes from archaea. Notably, 10% of the hits were for genes from a single organism, the soil archaeon Thorarchaeota strain OWC[26] (Supplementary Data 24).

***Rooting the sulfite oxidase family with minimal ancestor deviation***

The ancestor deviation was predicted for every branch of the sulfite oxidase phylogenetic tree. The five branches containing the slightest ancestor deviation were used to root the tree, and the topology was inspected for each. Each of the rooted trees supports a clade comprising the eukaryotic sulfite oxidase and nitrate reductase, and the Thaumarchaeota and Lutiacidiplasmatales putative sulfite oxidases to the exclusion of the other members of the family (Figure S2), indicating a common ancestry spanning domains of life.

***Duplication and loss of gene families originating in Lutiacidiplasmatales in comparison to ancestral gene families (Short name: Gene duplication)***

Increased duplication and loss rates were observed in laterally acquired gene families (in comparison to ancestral gene families) in Nitrososphaerales lineages[16]. In contrast, the gene families acquired by the Lutiacidiplasmatales LCA generally have lower rates of duplication and loss in Lutiacidiplasmatales lineages when compared to ancestral gene families (Supplementary Data 25). In Nitrososphaerales, these lineage-specific duplications and losses were theorised to have enabled these organisms to occupy refined niches within terrestrial and sediment environments. The lack of these changes in Lutiacidiplasmatales may indicate that the extant members of the order occupy a similar niche to their LCA. However, it is worth noting that all of the Lutiacidiplasmatales genomes in this study are from soil and that sequences with similarity to this group have been found in a myriad of non-soil environments.

Therefore, increased taxon sampling from different environments may reveal this order's gene content evolutionary history to be more complex than predicted here.

### Supplementary Figures

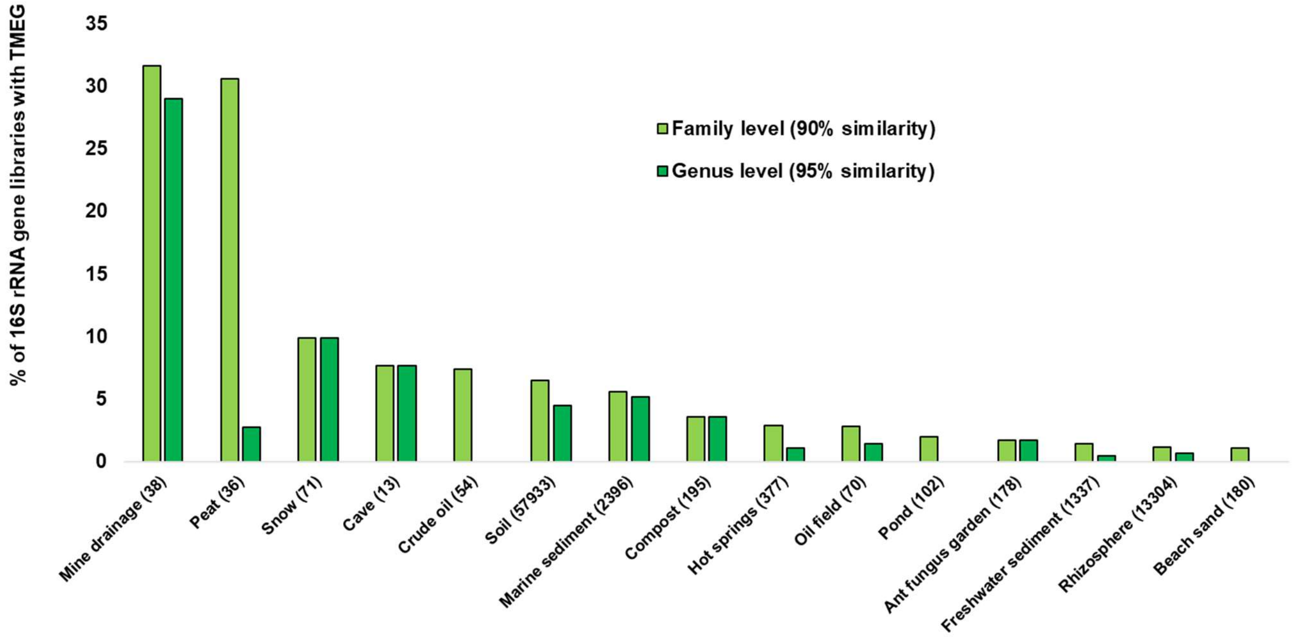


**Figure S1. Distribution of TMEG in publically available 16S rRNA gene libraries from many environments.** The 16S rRNA gene of AcS3-62 was queried against the extensive collection of 16S rRNA libraries in IMNGS[27] for reads of ≥ 400 bp that possessed ≥ 90 % sequence similarity.

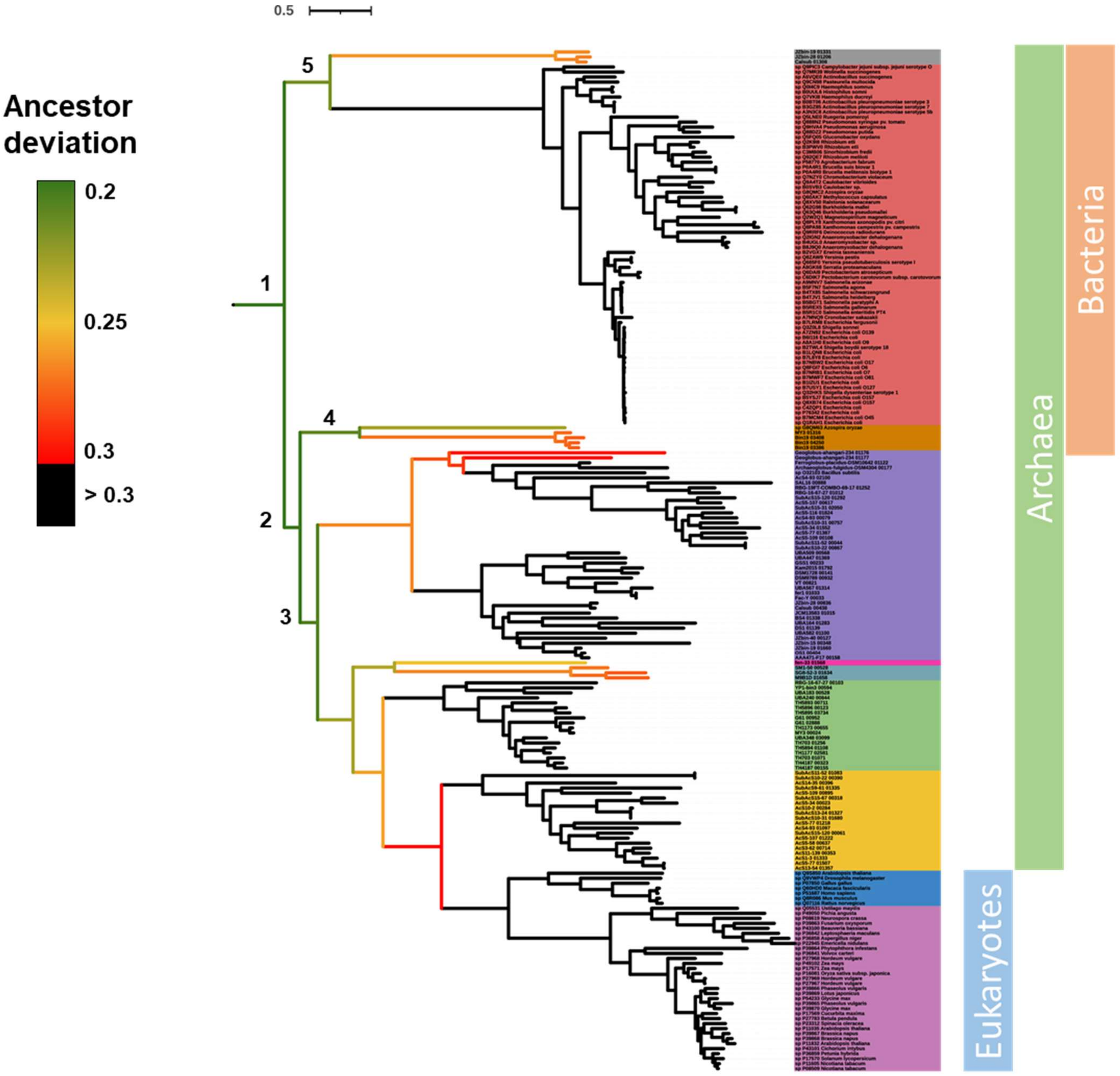


**Figure S2. Ancestor deviation in sulfite oxidase family tree.** Numbers indicate a rank of the most favoured roots by minimal ancestor deviation approach (**1** being the most favoured root). These roots show that the Lutiacidiplasmatales sulfite oxidases are the prokaryotic sequences most closely related to eukaryotic sulfite oxidases and nitrate reductases.

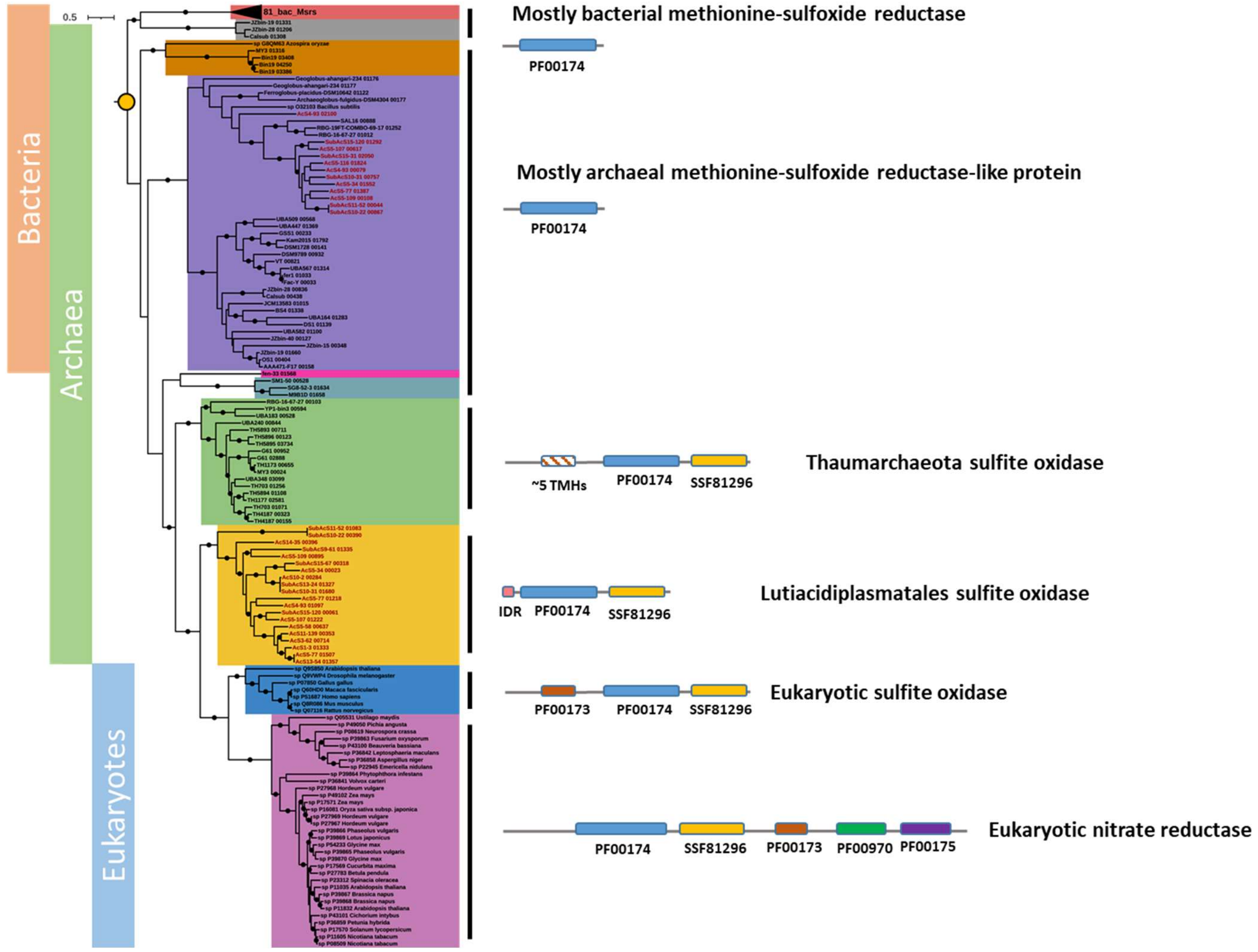


**Figure S3. Evolution of a putative archaeal class of sulfite oxidases.** Phylogenetic tree containing the sulfite oxidase superfamily members and a schematic of their conserved domains. Dots indicate branches with ≥ 70% of 2,000 UFBoot and 1,000 SH-aLRT replicates. Swiss-Prot sequences possessing the PF00174 domain and the previously identified Thaumarchaeota members of this family[16] were included in the analysis. The 81_bac_Msrs clade contains 81 bacterial methionine-sulfoxide reductase genes. Coloured bars on the left indicate the broad taxonomic affiliation of the protein clades. Overlapping of these bars indicates clades with proteins from more than one domain of life. Figures to the right of the tree are schematic representations of domain organisation in the corresponding protein clade with the following nomenclature: Transmembrane helices (TMHs) and Intrinsic disorder region (IDR), i.e. a natively unfolded region. The tree was rooted with minimal ancestor deviation (MAD), as described in Figure S2.

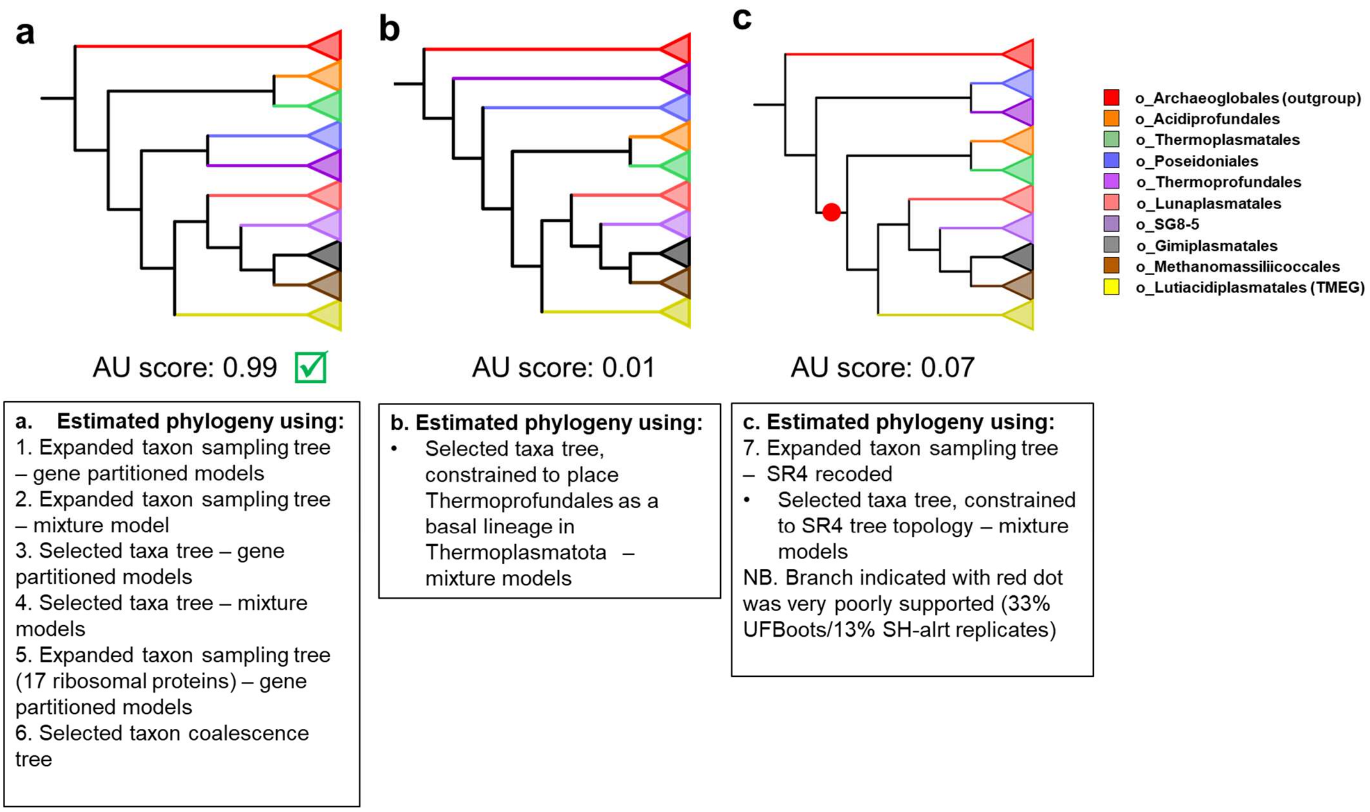


**Figure S4. Thermoplasmatota species tree estimation using multiple approaches and comparison of differing resulting topologies.** Approximately unbiased testing of the unconstrained selected taxa – gene partitioned models tree "**a**" and constrained trees with the topologies of "**b**" and "**c**" (AU score). Topology (a) received the highest likelihood and is the favoured hypothesis, although topology (c) could not be rejected at the $P < 0.05$ level. The red dot represents a poorly supported branch indicating an internal placement of Acidiprofundales and Thermoplasmatales in the Thermoplasmatota. Expanded taxon sampling trees are reconstructed from the full dataset of genomes, whereas the selected taxa trees are reconstructed from the high-quality dataset genomes.

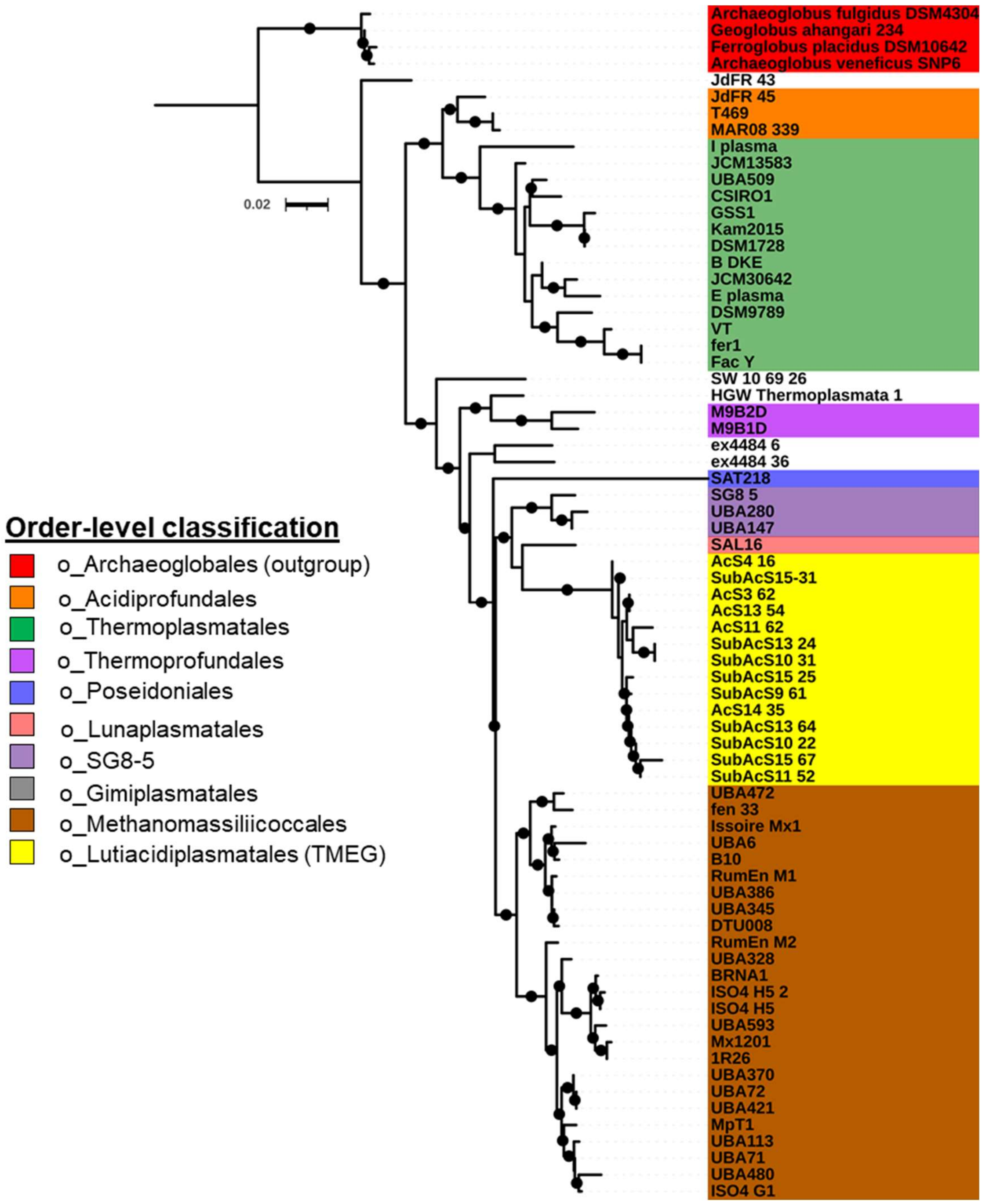


**Figure S5. Phylogenetic 16S rRNA gene tree of Thermoplasmatota genomes.** 16S rRNA gene sequences of ≥ 450 bp were extracted from the genomes analysed in this study. The final nucleotide alignment possessed 956 columns, 267 of which were parsimoniously informative. Dots indicate branches with ≥ 70% of 1,000 UFBoot replicates. The tree is rooted with the Archaeoglobales genes.

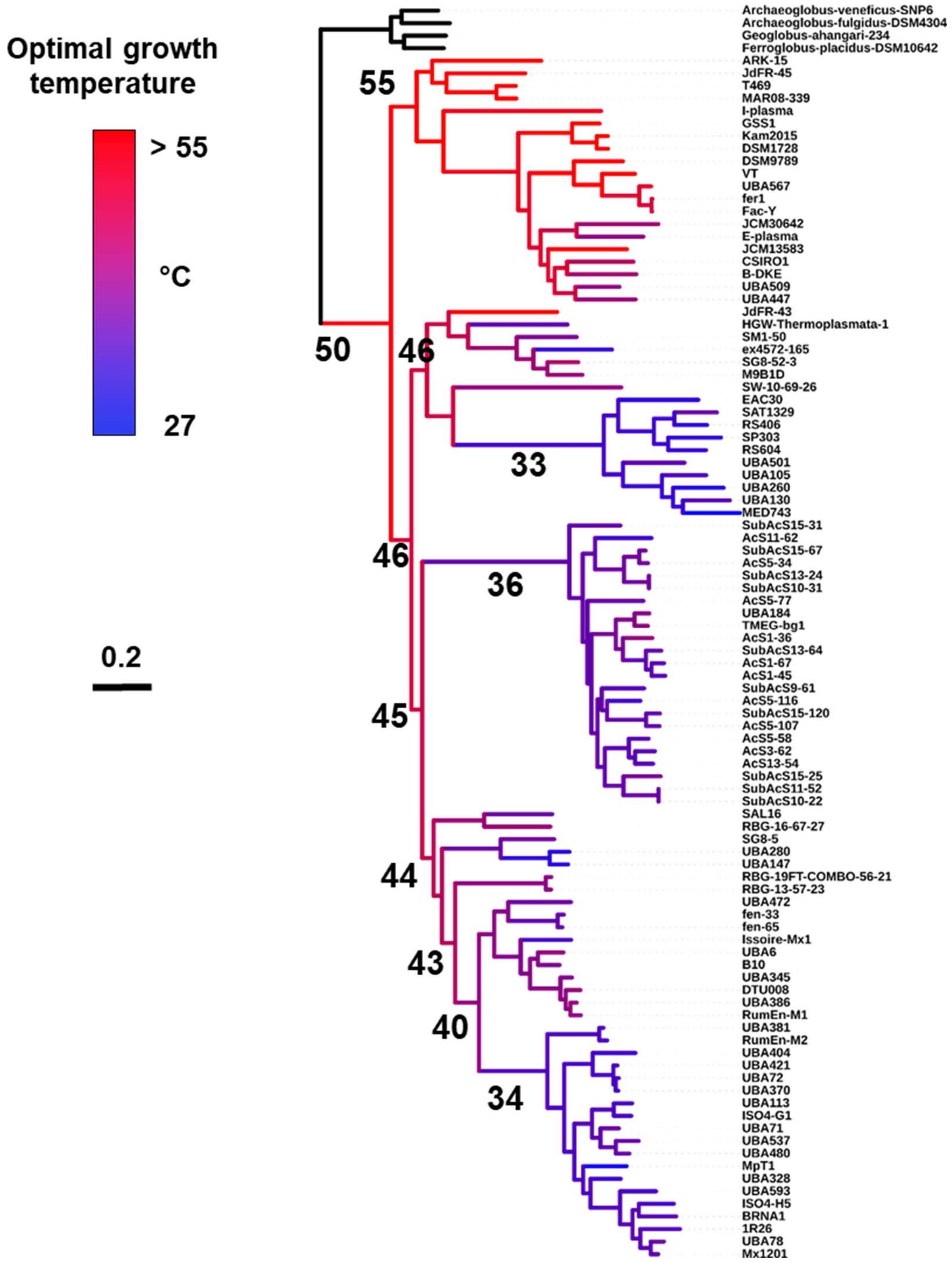


**Figure S6. Sequence-based prediction of thermal adaptation throughout Thermoplasmatota history.** Tome[28] was used to estimate the optimal growth temperatures (OGT) of extant Thermoplasmatota genomes. Ancestral OGTs were predicted based on a ridge regression approach[29]. Branches were coloured based on their predicted OGT, and key ancestral OGTs were specified on specific branches.

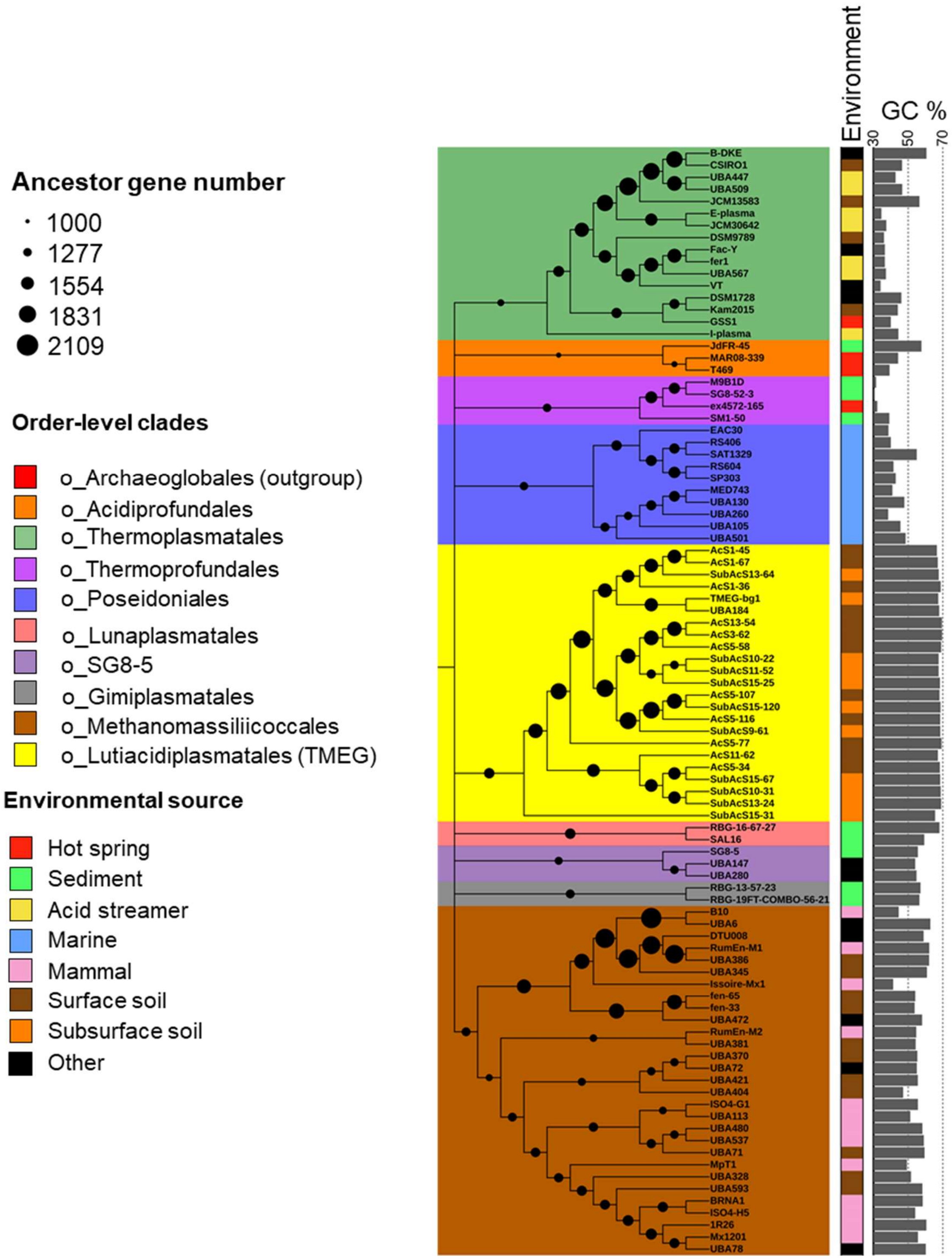


**Figure S7. Proteome size evolution and GC-content in Thermoplasmatota.** Colours across the phylogenetic tree indicate the order-level taxonomic affiliation of genomes. Dots sizes on branches represent the number of genes predicted in each ancestor reconstruction by gene tree –species tree reconciliation.

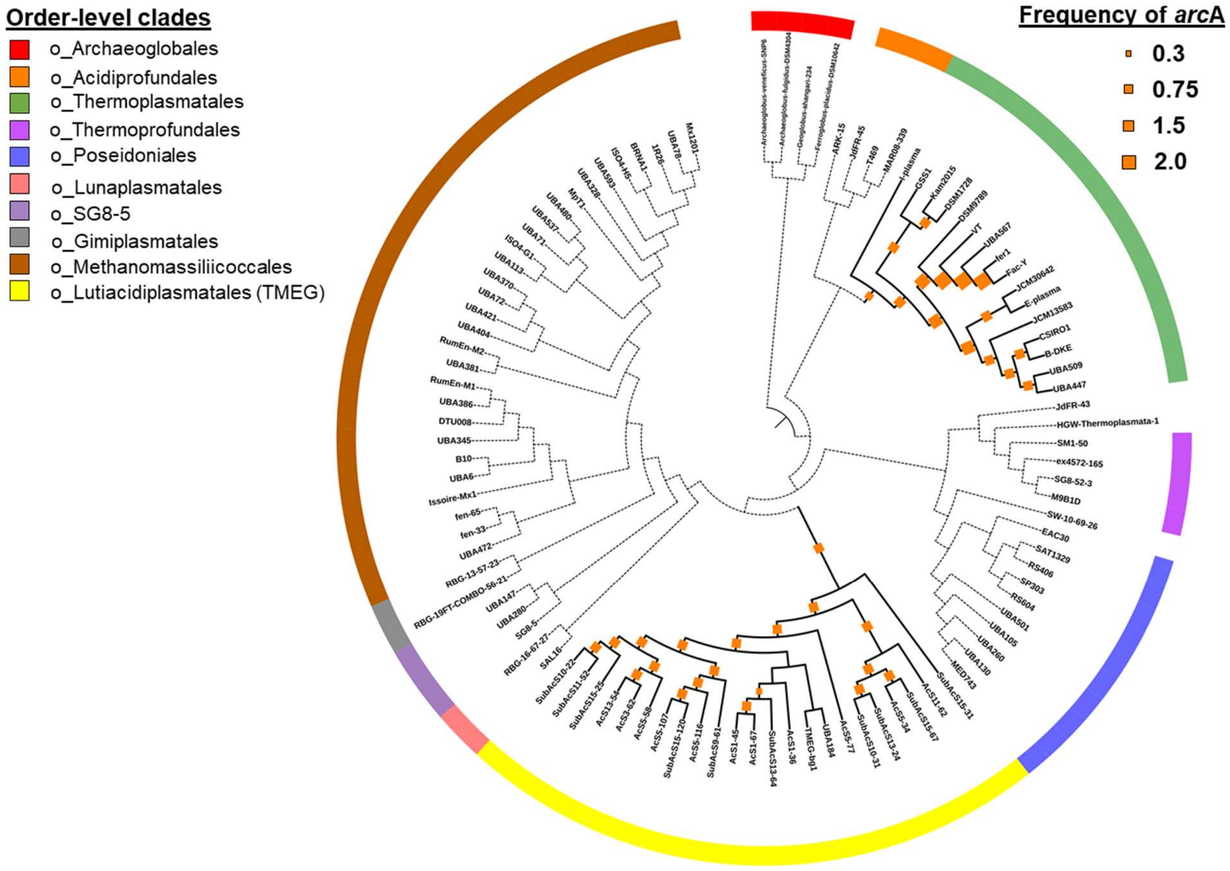


**Figure S8. Ancestral gene content comparison to detect multiple lateral acquisitions of the same gene family.** Gene tree-species tree reconciliation infers the frequency of a gene family on each branch of the species tree. Multiple lateral acquisitions (from inside or outside the phylum) were predicted when various lineages, but not the last common ancestor of these clades, possess a given gene family. In the example above, the arginine deiminase *arc*A gene family is present in the Lutiacidiplasmatales and Thermoplasmatales lineages but is not present in Thermoplasmatales last common ancestor. This indicates *arc*A entered these orders by separate lateral gene transfers rather than vertical inheritance from a common ancestor. Individual gene trees with a diverse representation of *arc*A genes from multiple phyla can then be used to infer whether both lateral gene transfers were from different phyla or whether *arc*A was transferred between the Lutiacidiplasmatales and Thermoplasmatales.

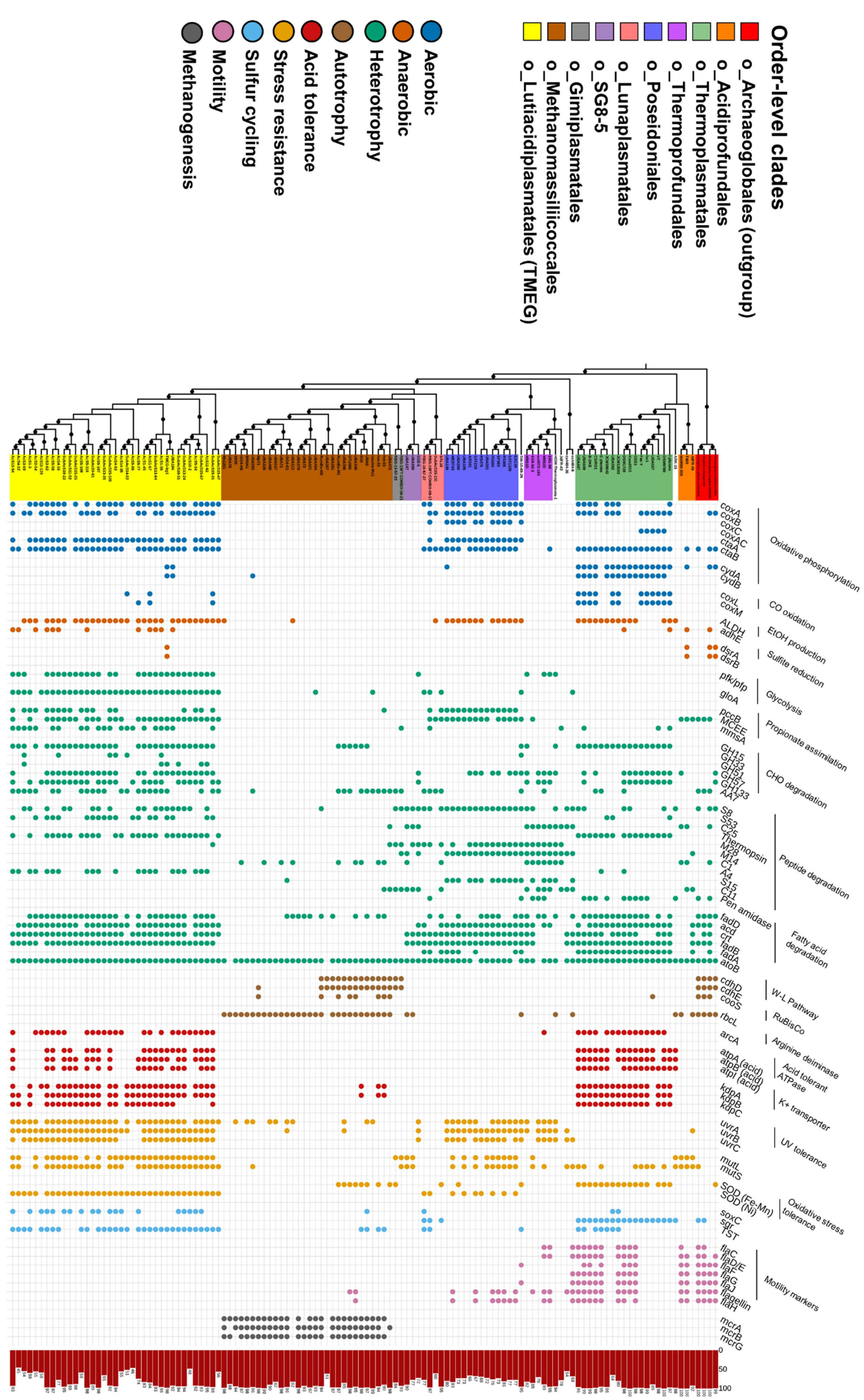
Order-level clades
o_Archaeoglobales (outgroup)
o_Acidiprofundales
o_Thermoplasmatales
o_Thermoprofundales
o_Poseidoniales
o_Lunaplasmatales
o_SG8-5
o_Gimiplasmatales
o_Methanomassiliicoccales
o_Lutiacidiplasmatales (TMEG)
Aerobic
Anaerobic
Heterotrophy
Autotrophy
Acid tolerance
Stress resistance
Sulfur cycling
Motility
Methanogenesis
coxA
coxB
coxC
coxAC
ctaA
ctaB
cydA
cydB
Oxidative phosphorylation
coxL
coxM
CO oxidation
ALDH
adhE
EtOH production
dsrA
dsrB
Sulfite reduction
pfk/pfp
gloA
Glycolysis
pccB
MCEE
mmsA
Propionate assimilation
GH15
GH33
GH51
GH57
GH133
AA7
CHO degradation
S8
S53
C25
Thermopsin
M28
M14
C1
A4
S15
C11
Pen amidase
Peptide degradation
fadD
acd
crt
fadB
fadA
atoB
Fatty acid degradation
cdhD
cdhE
cooS
W-L Pathway
rbcL
RuBisCo
arcA
Arginine deiminase
atpA (acid)
atpB (acid)
atpI (acid)
Acid tolerant ATPase
kdpA
kdpB
kdpC
K+ transporter
uvrA
uvrB
uvrC
UV tolerance
mutL
mutS
SOD (Fe-Mn)
SOD (Ni)
Oxidative stress tolerance
soxC
sqr
TST
flaC
flaD/E
flaF
flaG
flaJ
flagellin
flaH
Motility markers
mcrA
mcrB
mcrG
0
50
100

**Figure S9. Metabolism of the Thermoplasmatota.** The presence of selected genes was assessed for all genomes in the analysis. *cox*A (heme-copper oxygen reductase, subunit A; K02274), *cox*B (heme-copper oxygen reductase, subunit B; K02275), *cox*C (heme-copper oxygen reductase, subunit C; PF00510), *cox*AC (heme-copper oxygen reductase, subunit AC; K15408), *cta*A (heme a synthase; K02259), *cta*B (heme o synthase; K02257), *cyd*A (cytochrome bd ubiquinol oxidase, subunit A; K00425), *cyd*B (cytochrome bd ubiquinol oxidase, subunit B; K00426), *cox*L (aerobic carbon-monoxide dehydrogenase, large subunit; K03520), *cox*M (aerobic carbon-monoxide dehydrogenase, medium subunit; K03519), ALDH (aldehyde dehydrogenase (NAD+); K00128), *adh*E (alcohol dehydrogenase; K0407), *dsr*AB (dissimilatory sulfite reductase, subunits A and B; K11180 and K11181), *pfk/pfp* (ATP-dependent phosphofructokinase; K21071), *glo*A (lactoylglutathione lyase; K01759), *pcc*B (propionyl-CoA carboxylase, beta chain, K01966), MCEE (methylmalonyl-CoA/ethylmalonyl-CoA epimerase; K05606), *mms*A (methylmalonate-semialdehyde dehydrogenase; K00140), GH# (glycoside hydrolase family #; dbCAN), Pen amidase (Penicillin amidase), *fad*D (long-chain acyl-CoA synthetase; K01897), *acd* (acyl-CoA dehydrogenase; K00249), *crt* (enoyl-CoA hydratase; K01715), *fad*B (3-hydroxybutyryl-CoA dehydrogenase; K00074), *fad*A (acetyl-CoA acyltransferase; K00632), *ato*B (acetyl-CoA C-acetyltransferase; K00626), *cdh*DE (acetyl-CoA decarbonylase/synthase complex D; K00194 and E; K00197 subunits), *coo*S (anaerobic carbon-monoxide dehydrogenase catalytic subunit; K00198), *rbc*L (ribulose-bisphosphate carboxylase large chain; K01601), *arc*A (arginine deiminase; K01478), *atp*ABI (acid) (V/A-type atpase A, B and I subunits; K02117, K02118 and K02123, plus gene tree analysis), *kdp*ABC (K+ transporting ATPase subunits A, B and C; K01546, K01547 and K01548), *uvr*ABC (excinuclease subunits A, B and C; K03701, K03702 and K03703), *mut*LS (DNA mismatch repair proteins L and S; K03572 and K03555), SOD (Fe-Mn) (superoxide dismutase Fe-Mn family; K04564), SOD (Ni) (nickel superoxide dismutase; K00518), *sox*C (sulfite oxidase; TIGR04555), *sqr* (sulfide:quinone oxidoreductase; K17218), TST (thiosulfate/3-mercaptopyruvate sulfurtransferase; K01011), *fla*# (archaellum subunits C; arCOG05119, D/E ; arCOG02964, F; arCOG01824, G; arCOG01822 and J; arCOG01809), flagellin (archaeal flagellin; PF01917) and *fla*H (archaellum subunits C;PF06745) and *mrc*ABG (methyl-coenzyme M reductase A; K00399, B; K00401, C; K00402 subunits). The predicted completeness of each genome sequence is indicated in the far right red bar chart.

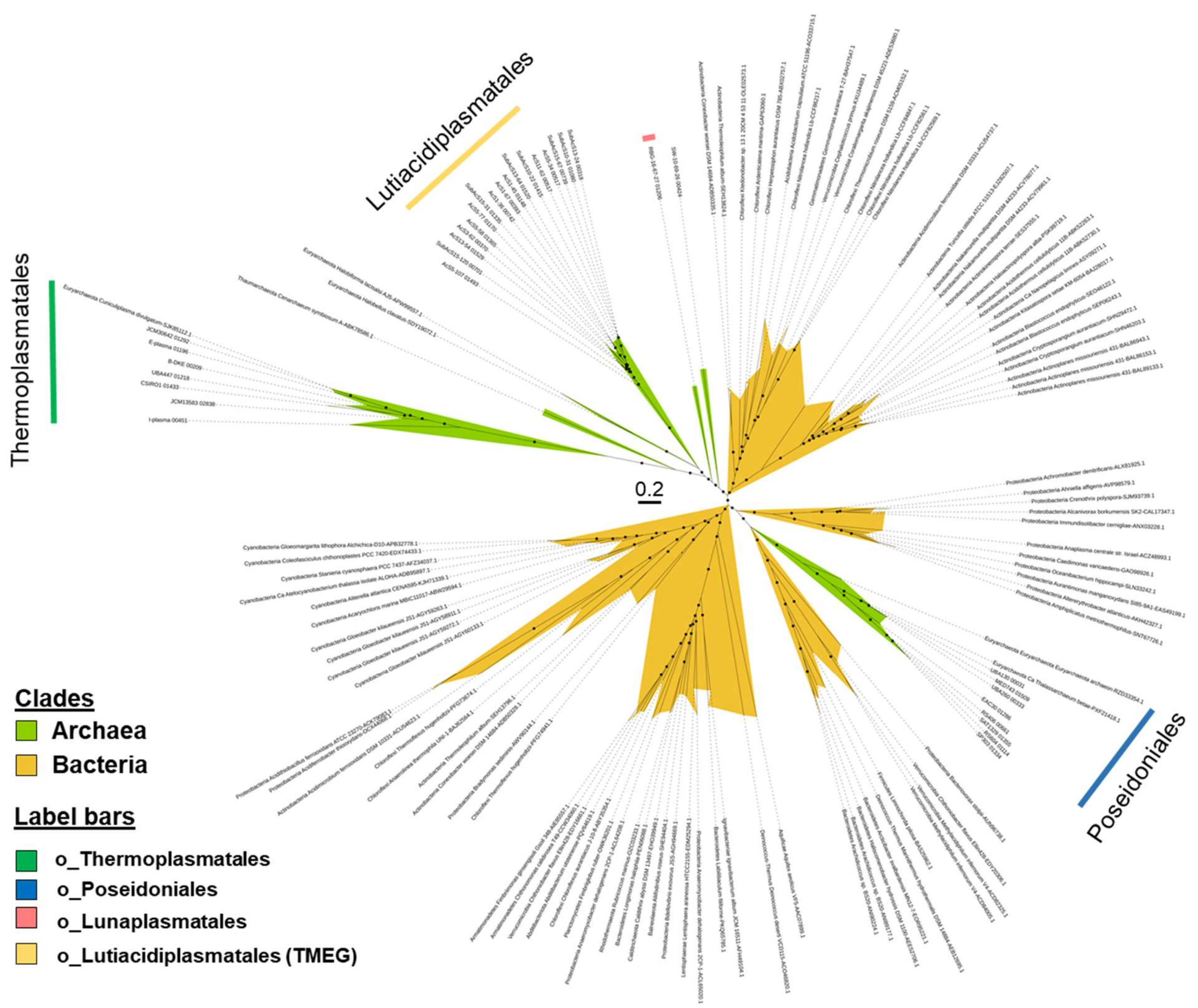


**Figure S10. Phylogeny of the heme-copper oxygen reductase subunit A (*cox*A) gene.** Dots indicate branches with ≥ 70% of 1,000 UFBoot replicates. The tree was estimated using the model LG+F+R7.

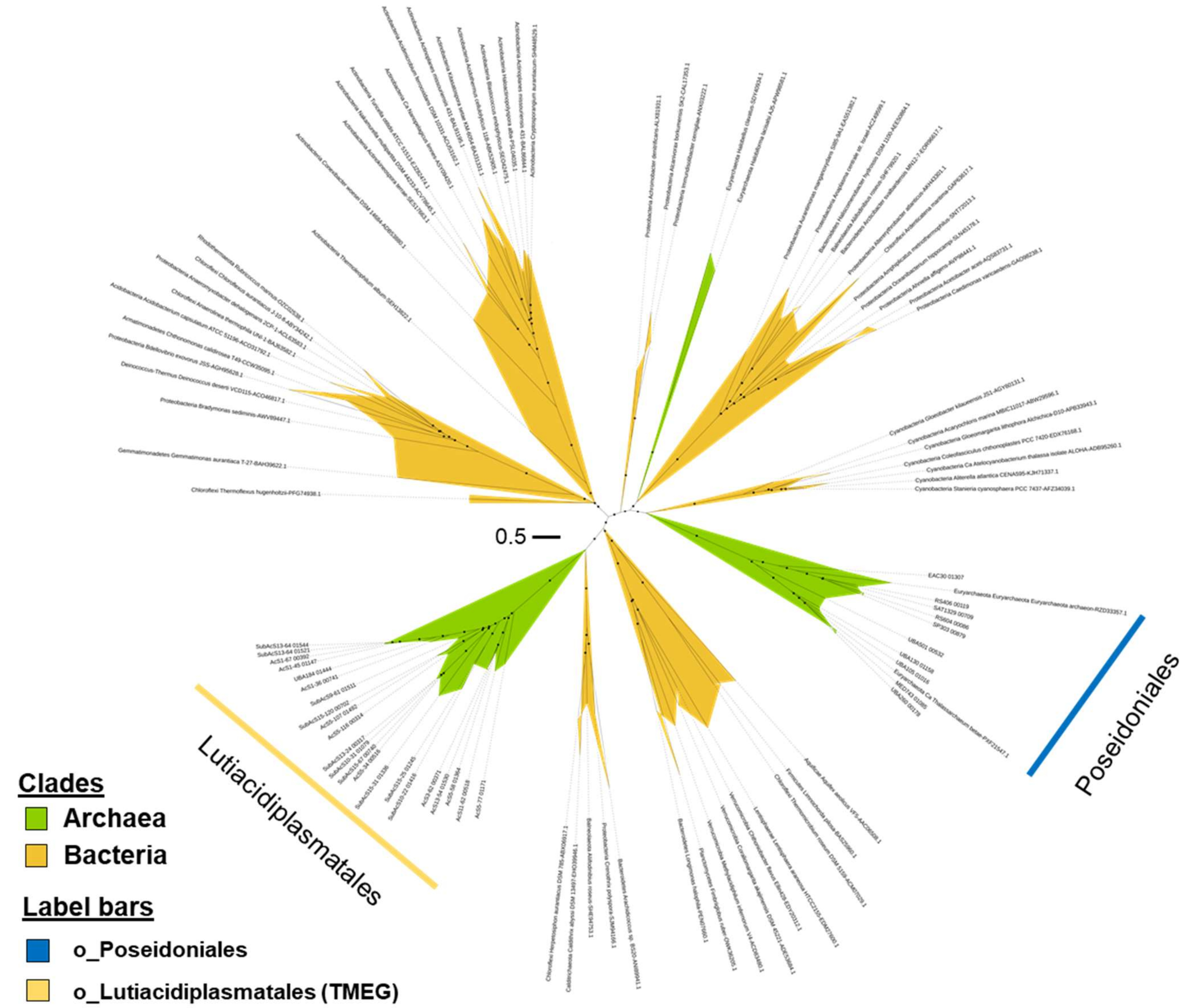


**Figure S11. Phylogeny of the heme A synthase (*cta*A) gene.** Dots indicate branches with ≥ 70% of 1,000 UFBoot replicates. The tree was estimated using the model LG+F+R7.

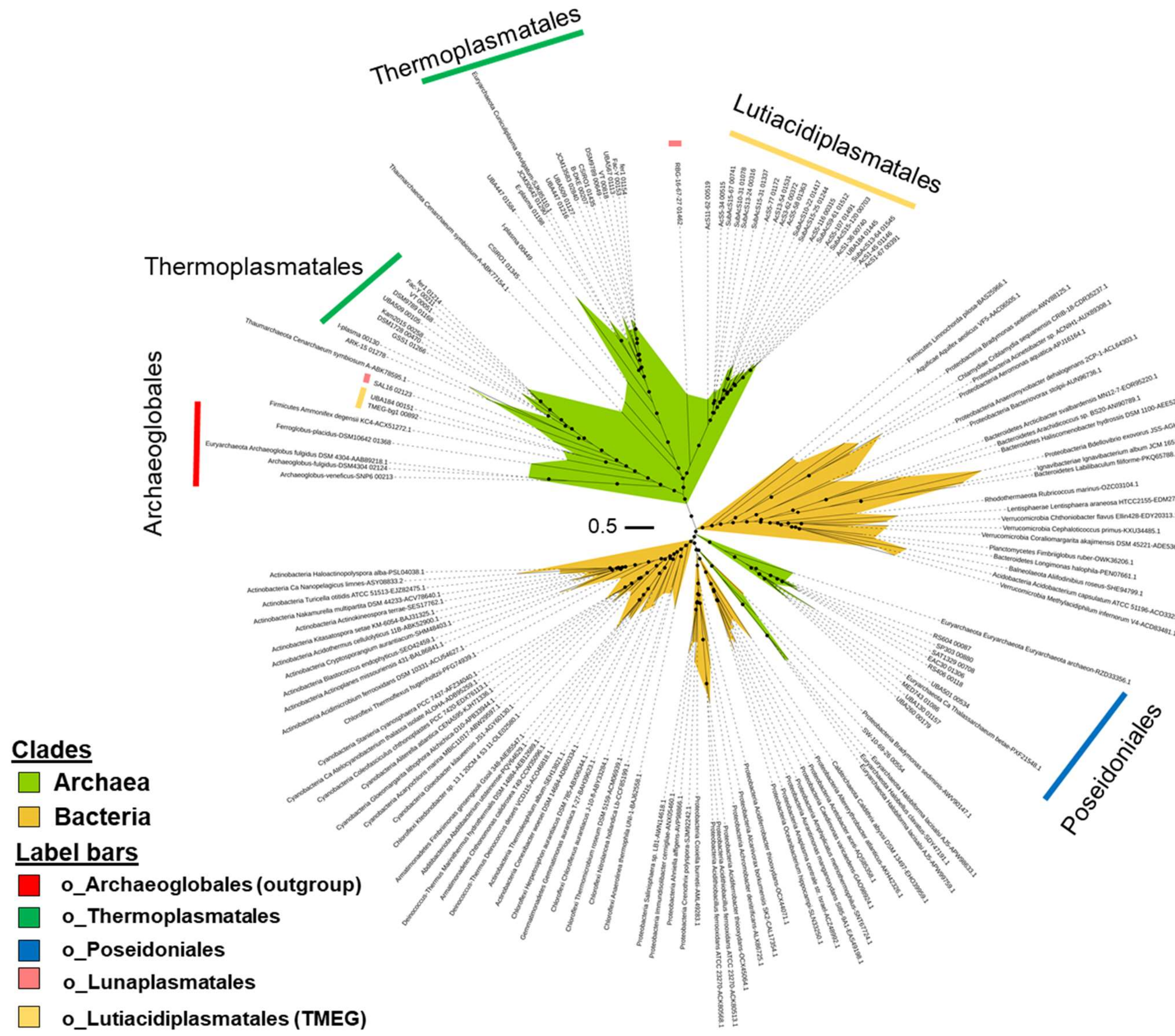


**Figure S12. Phylogeny of the protoheme IX farnesyltransferase (*cta*B) gene.** Dots indicate branches with ≥ 70% of 1,000 UFBoot replicates. The tree was estimated using the model LG+F+R7.

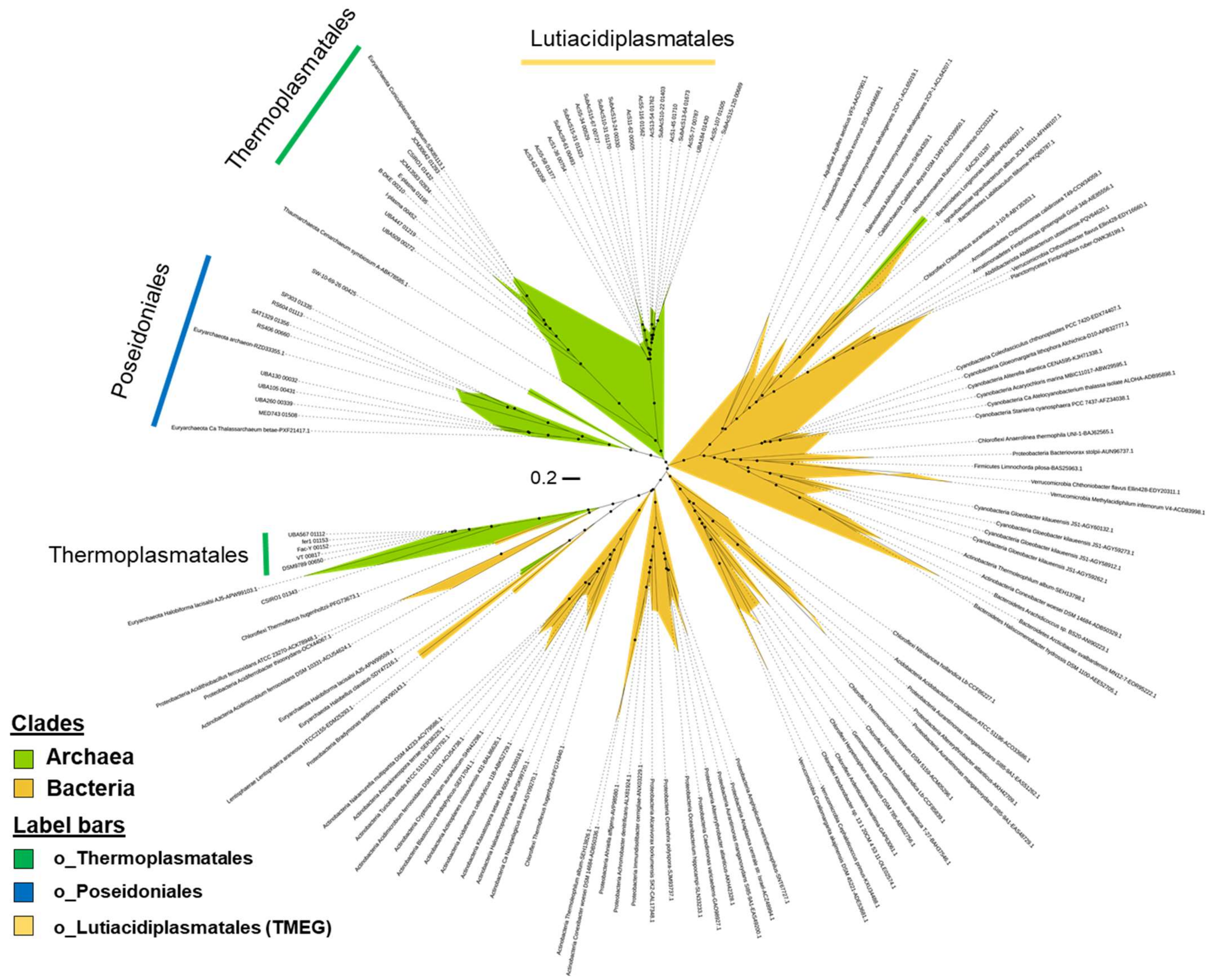


**Figure S13. Phylogeny of the heme-copper oxygen reductase subunit B (*cox*B) gene.** Dots indicate branches with ≥ 70% of 1,000 UFBoot replicates. The tree was estimated using the model LG+R6.

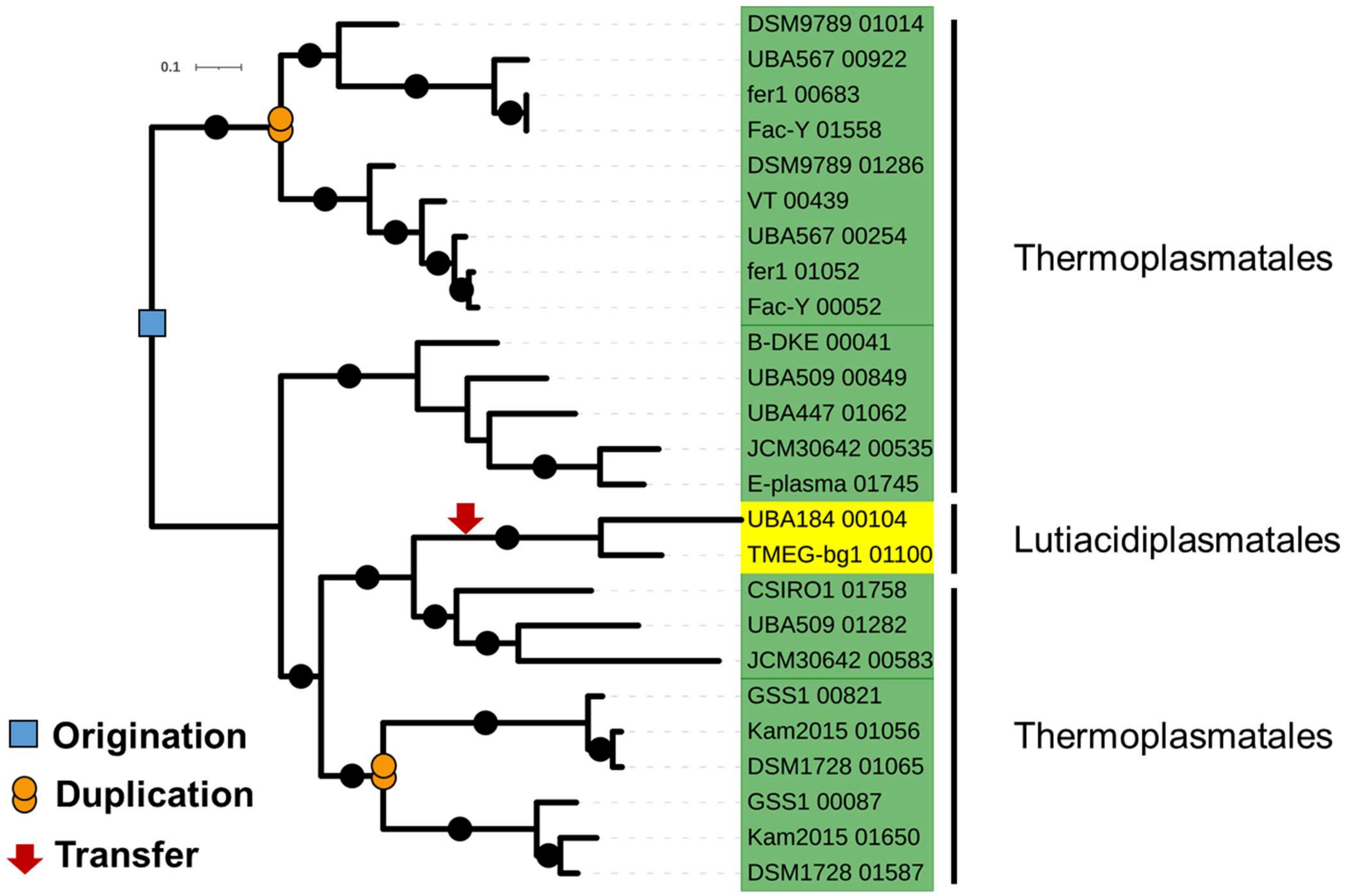


**Figure S14. Evolutionary history of the cytochrome bd ubiquinol oxidase (*cyd*A) gene in Thermoplasmatota.** The *cyd*A gene is predicted to have been acquired by lateral gene transfer by a Thermoplasmatales ancestor. The gene was subsequently duplicated within the Thermoplasmatales and laterally transferred to the LCA of Lutiacidiplasmatales genomes, UBA184 and TMEG-bg1. The *cyd*A gene phylogenetic tree was estimated using the LG+F+G4 model and rooted using minimal ancestor deviation (MAD). Dots indicate branches with ≥ 70% of 1,000 UFBoot replicates.

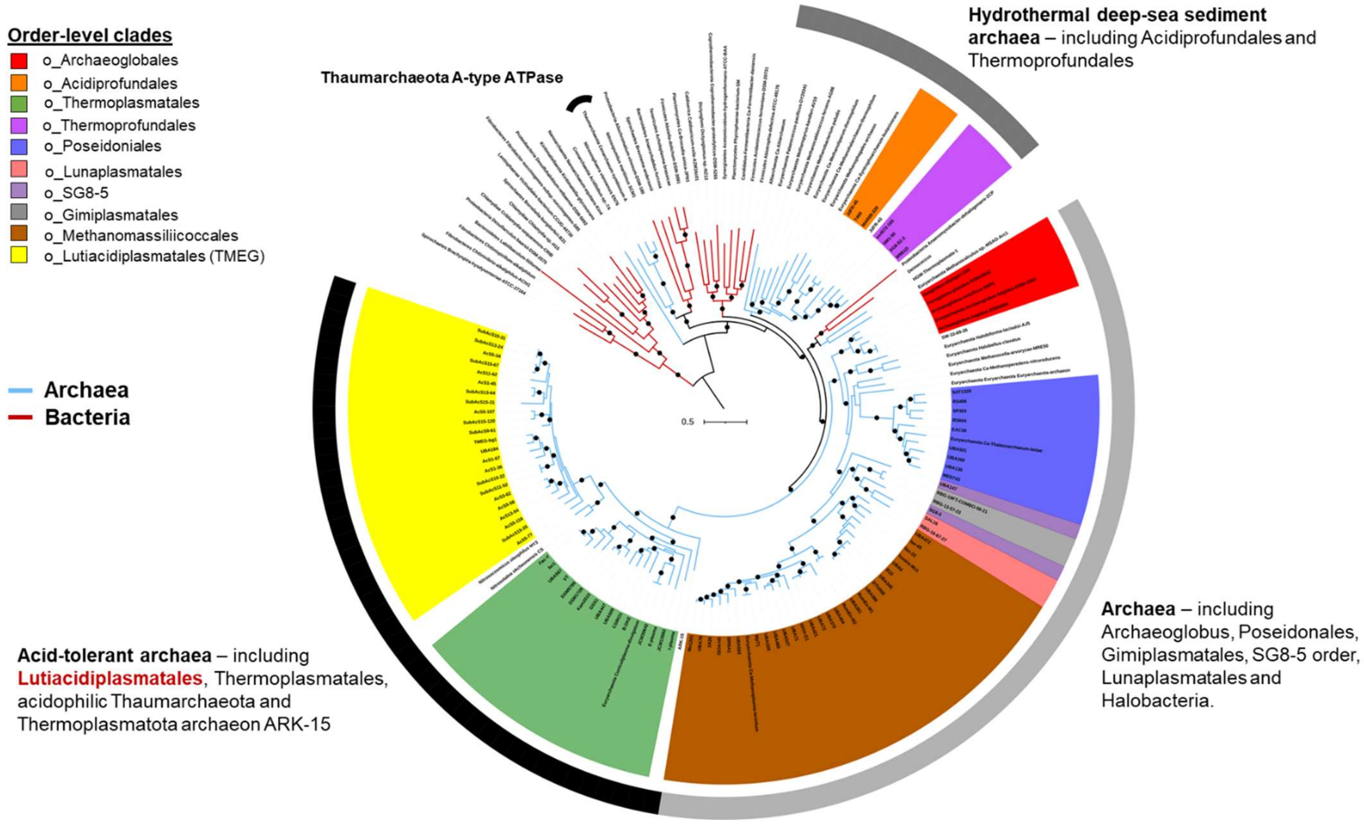


**Figure S15. Phylogeny of the V/A-ATPase.** Lutiacidiplasmatales cluster with the acid-tolerant archaea. The three largest subunits of V/A-ATPase (*atp*A, B and I) were individually aligned and then concatenated into a single partitioned supermatrix. A supermatrix tree was then estimated using the best fitting model for each partition and rooted using minimal ancestor deviation (MAD). Dots indicate branches with ≥ 70% of 1,000 UFBoot replicates.

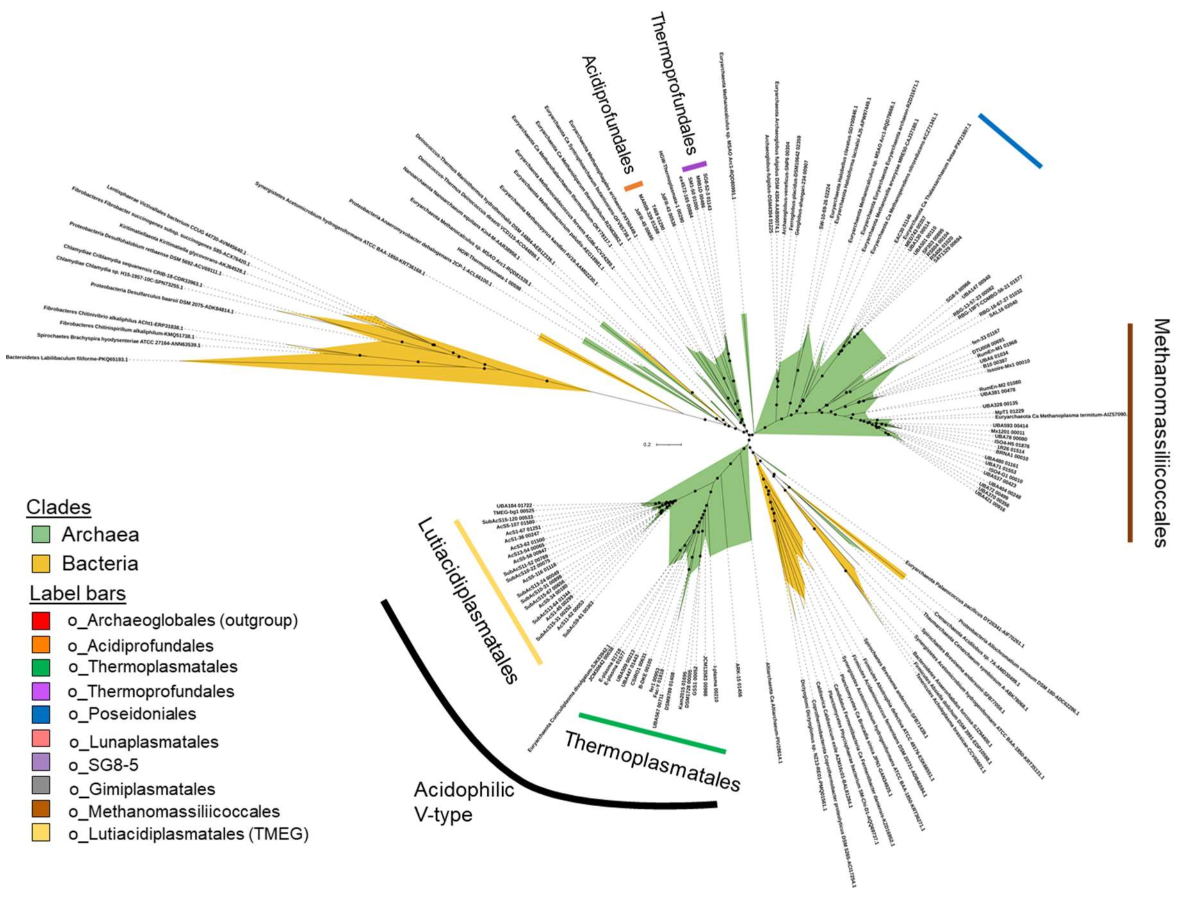


**Figure 16. Phylogeny of the V/A-ATPase subunit A (*atp*A) gene.** Dots indicate branches with ≥ 70% of 1,000 UFBoot replicates. The tree was estimated using the model LG+R6.

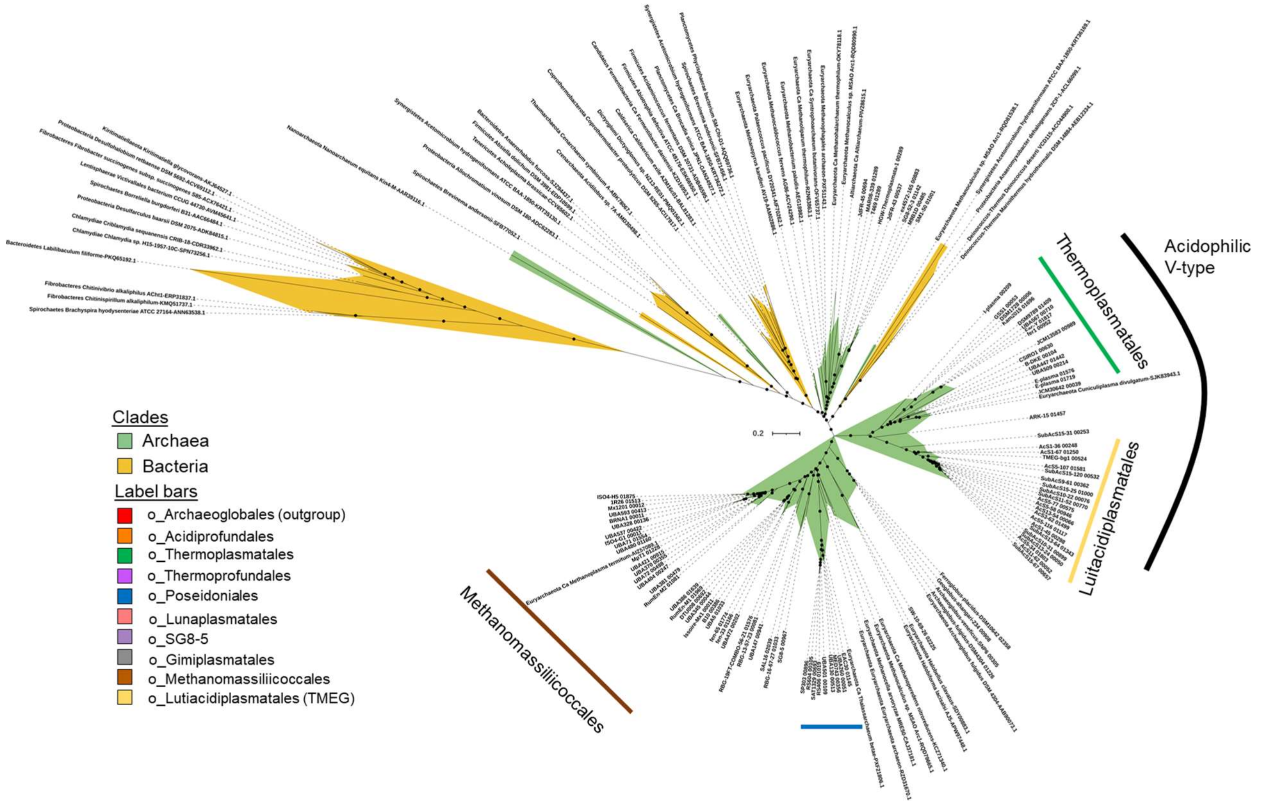


**Figure 17. Phylogeny of the V/A-ATPase subunit B (*atp*B) gene.** Dots indicate branches with ≥ 70% of 1,000 UFBoot replicates. The tree was estimated using the model LG+R6.

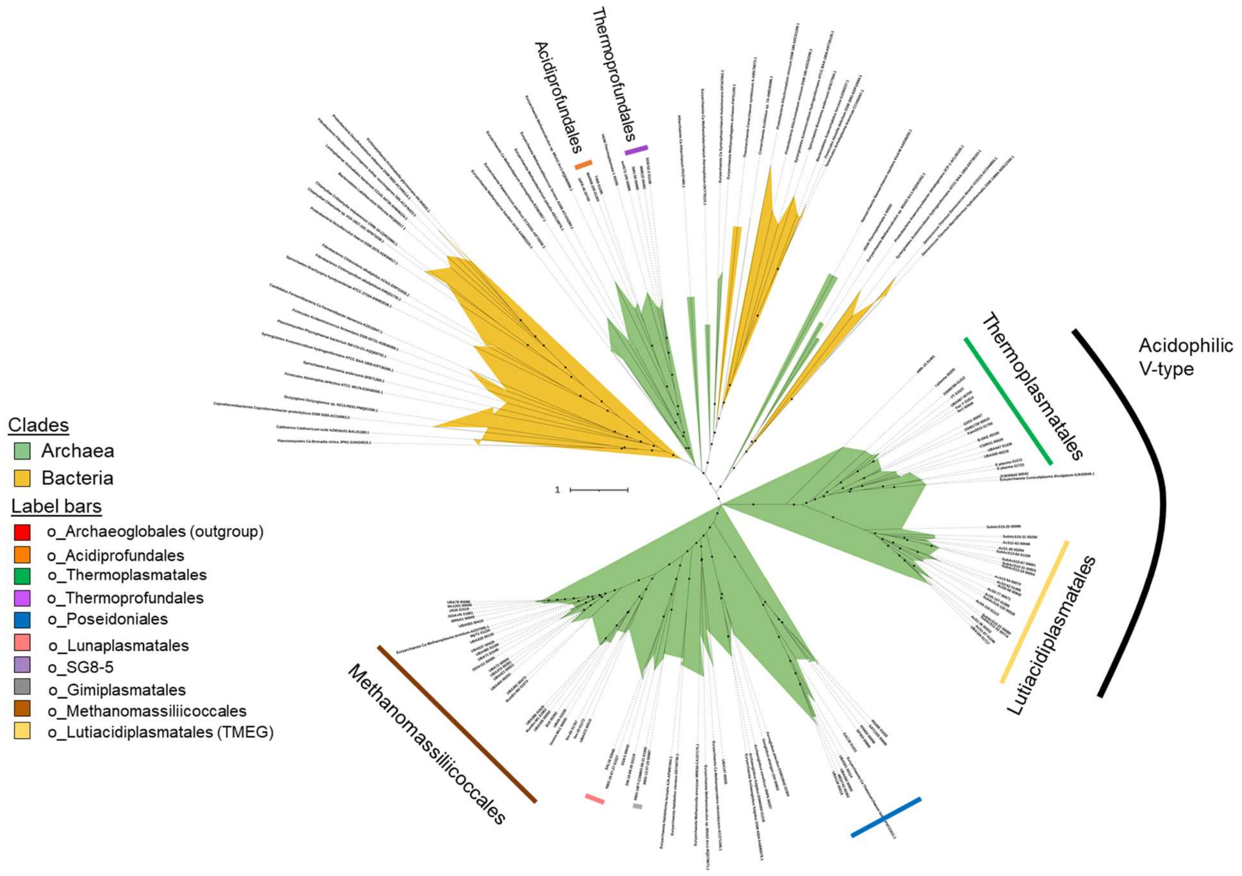


**Figure 18. Phylogeny of the V/A-ATPase subunit I (*atp*I) gene.** Dots indicate branches with ≥ 70% of 1,000 UFBoot replicates. The tree was estimated using the model LG+F+R7.

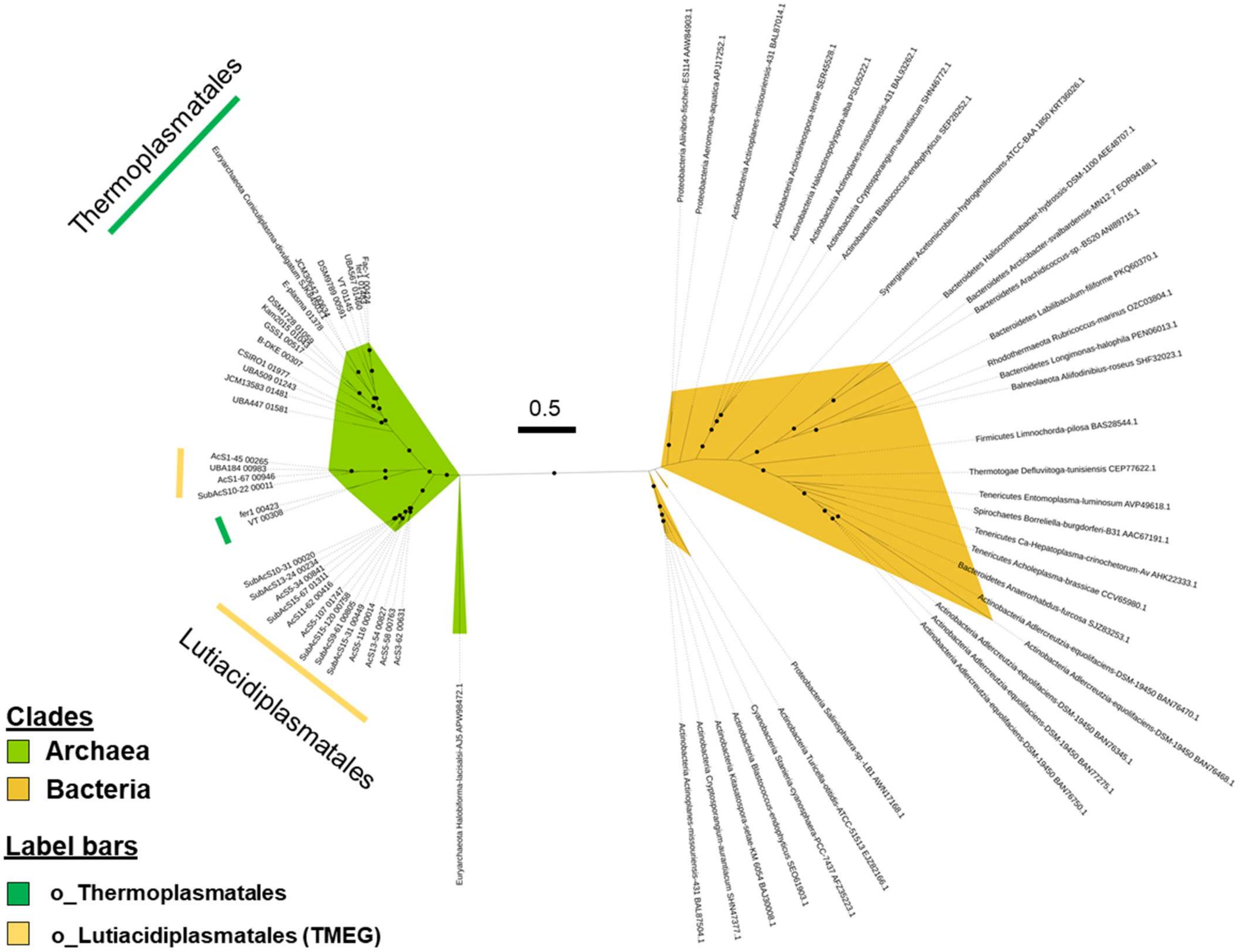


**Figure 19. Phylogeny of the arginine deiminase (*arc*A) gene.** Dots indicate branches with ≥ 70% of 1,000 UFBoot replicates. The tree was estimated using the mode LG+R5.

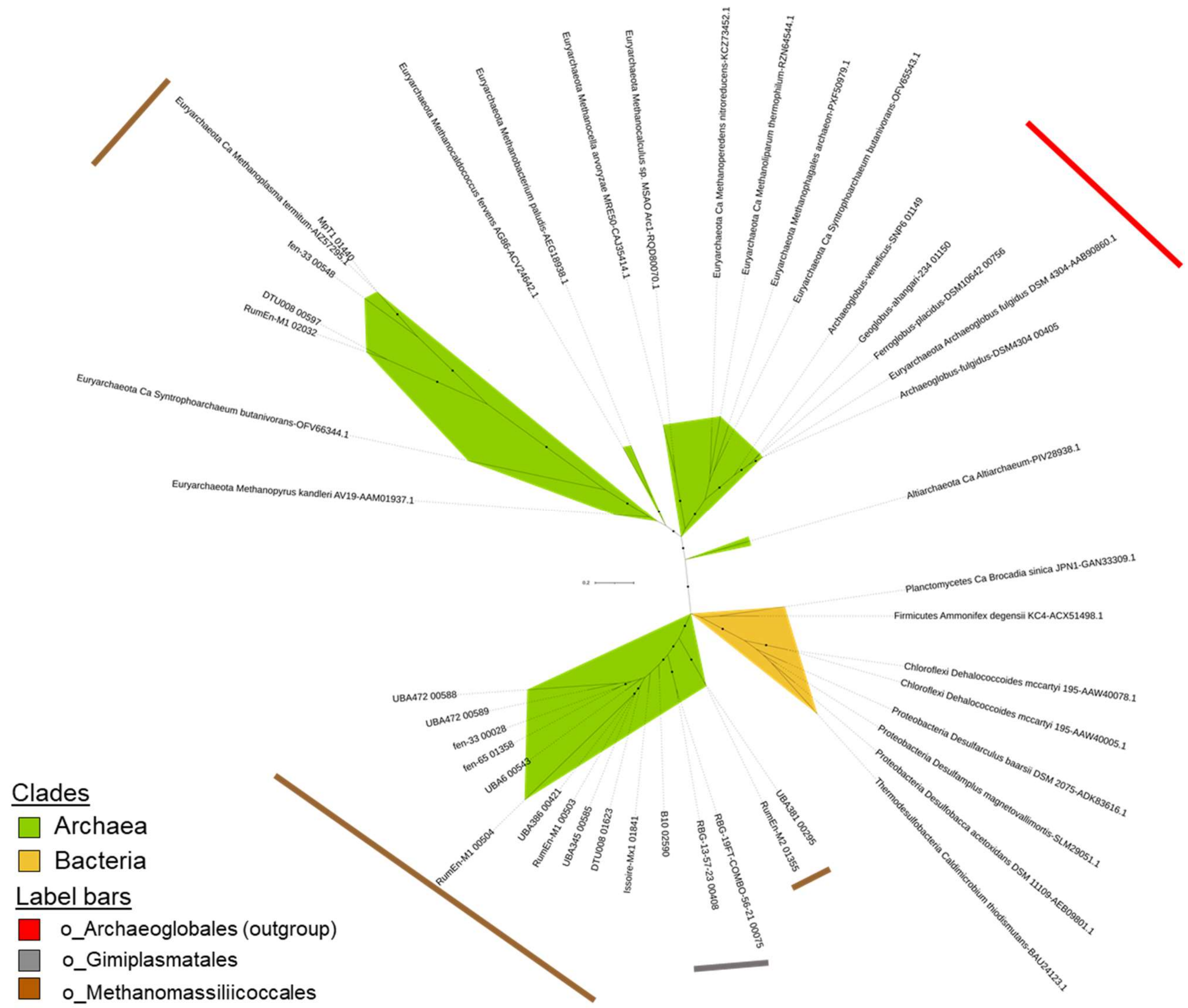


**Figure S20. Phylogeny of the CO dehydrogenase/acetyl-CoA synthase complex subunit gamma (*cdh*E).** Dots indicate branches with ≥ 70% of 1,000 UFBoot replicates. The tree was estimated using the model LG+I+G4.

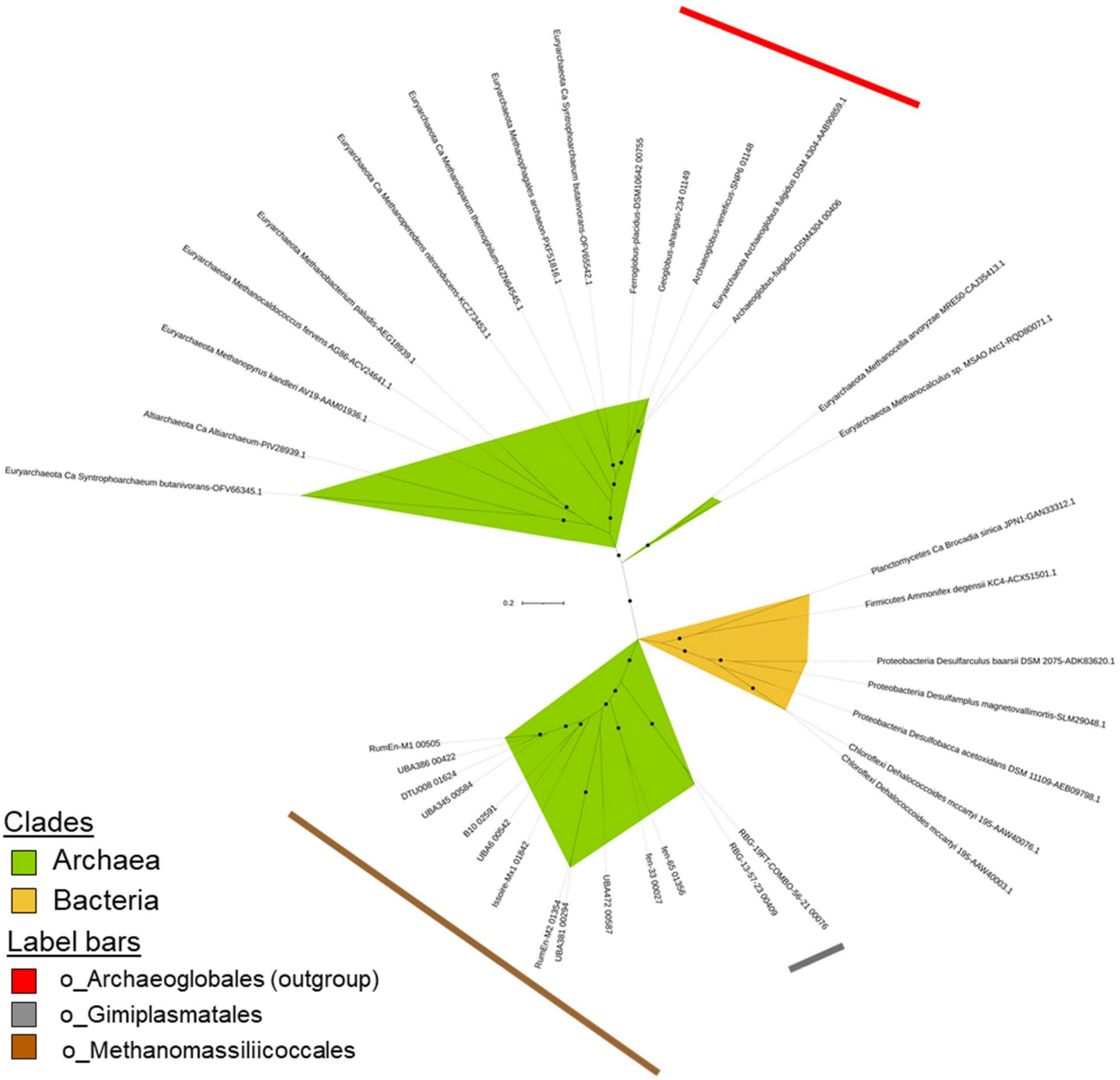


**Figure S21. Phylogeny of the CO dehydrogenase/acetyl-CoA synthase complex subunit delta (*cdh*D).** Dots indicate branches with ≥ 70% of 1,000 UFBoot replicates. The tree was estimated using the model LG+I+G4.

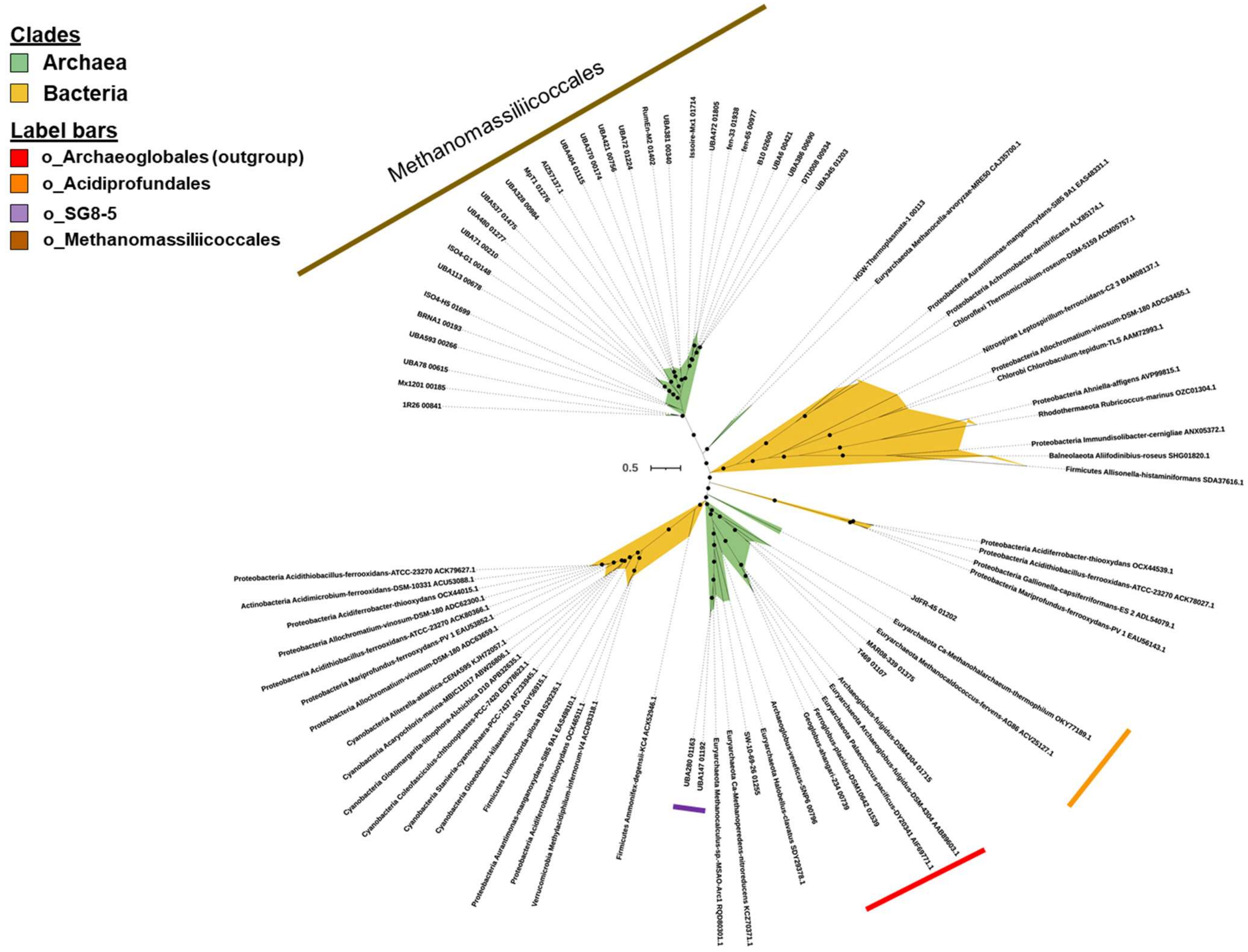


**Figure S22. Phylogeny of the ribulose bisphosphate carboxylase, large chain subunit (*rbc*L).** Dots indicate branches with ≥ 70% of 1,000 UFBoot replicates. The tree was estimated using the model LG+R5.

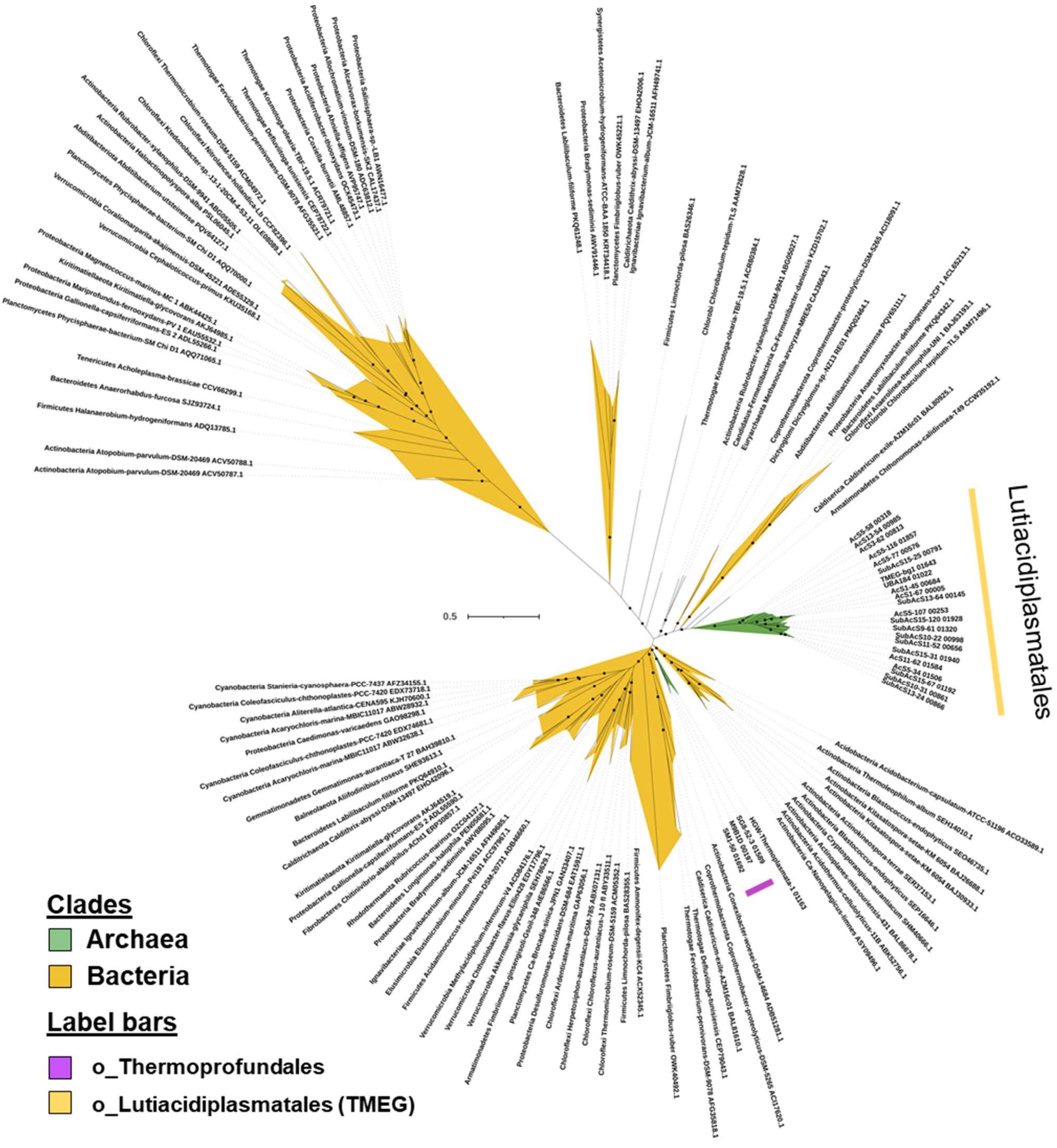


**Figure S23. Phylogeny of the ATP-dependent phosphofructokinase (*pfk*).** Dots indicate branches with ≥ 70% of 1,000 UFBoot replicates. The tree was estimated using the model LG+R6.

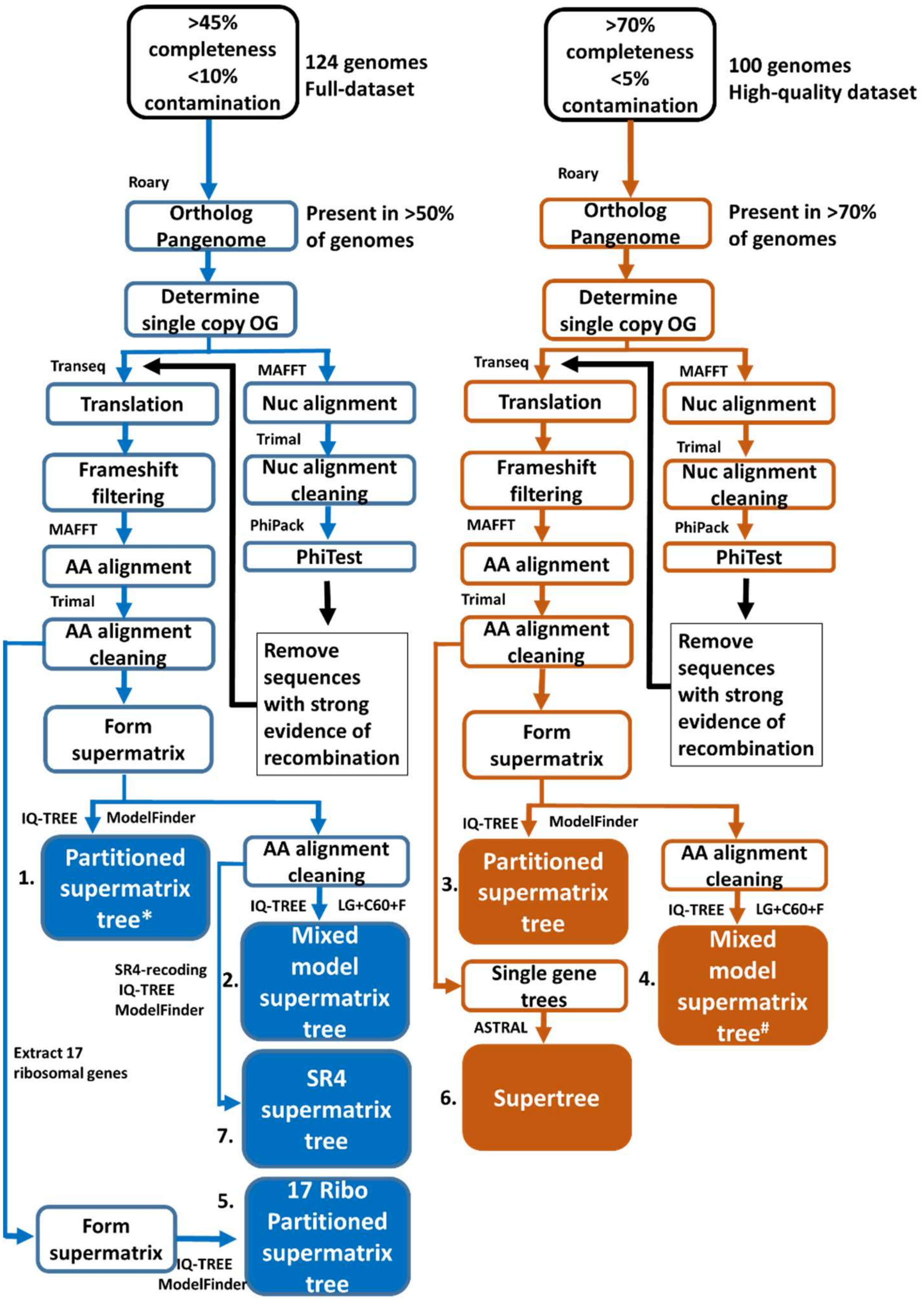


**Figure S24. Schematic workflow of construction of six phylogenomic trees estimating Thermoplasmatota evolutionary relationships.** Tools used in the blue and orange branches of the workflow are shown next to their stage of use. The numbered colour filled boxes indicate the six trees created in this workflow. The asterisk (*) indicates the tree used in the creation of Figure 2, whereas the hashtag (#) indicates the tree used as a species tree in the gene tree-species tree reconciliation. Numbers on trees correspond to those in Figure S4.

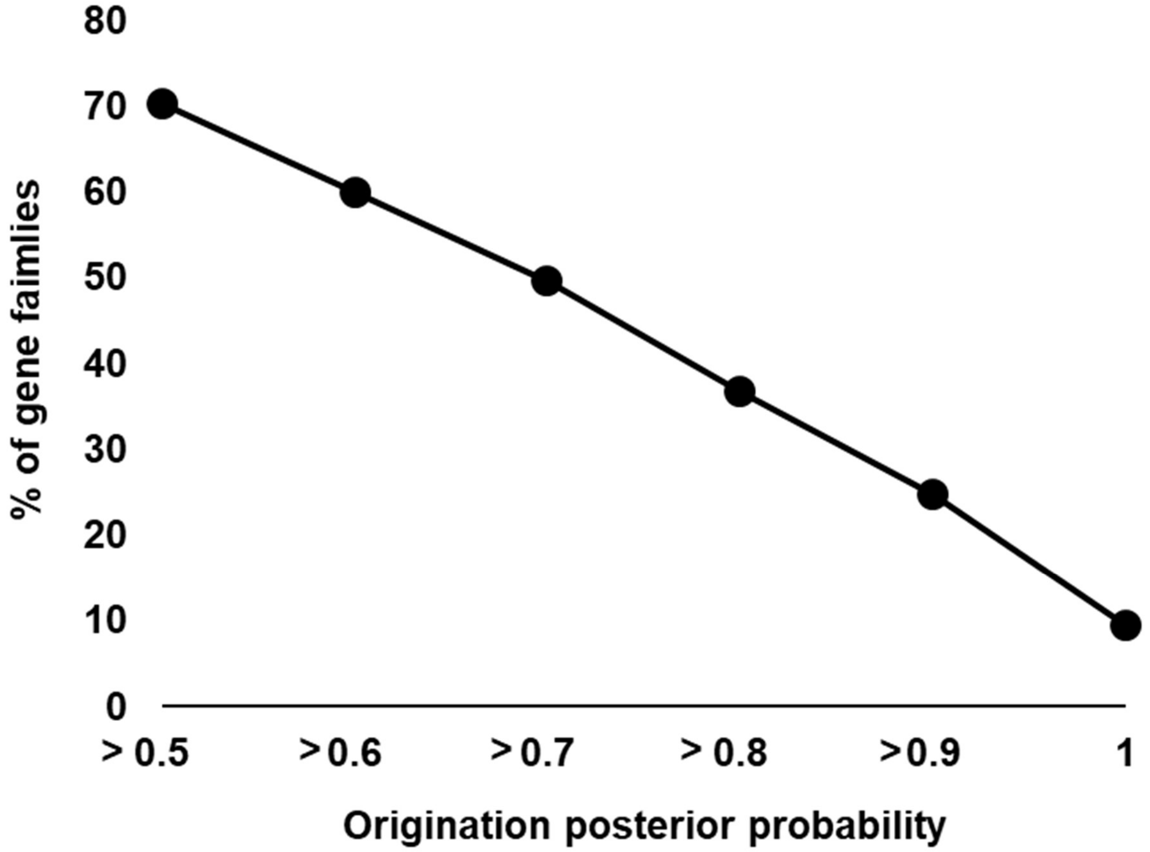


**Figure S25. Percentage of the 6,050 gene families predicted to have a single point of origination against increasing stringency of the posterior probability.** There is a linear decrease in the percentage of gene families with increasing origination posterior probability.

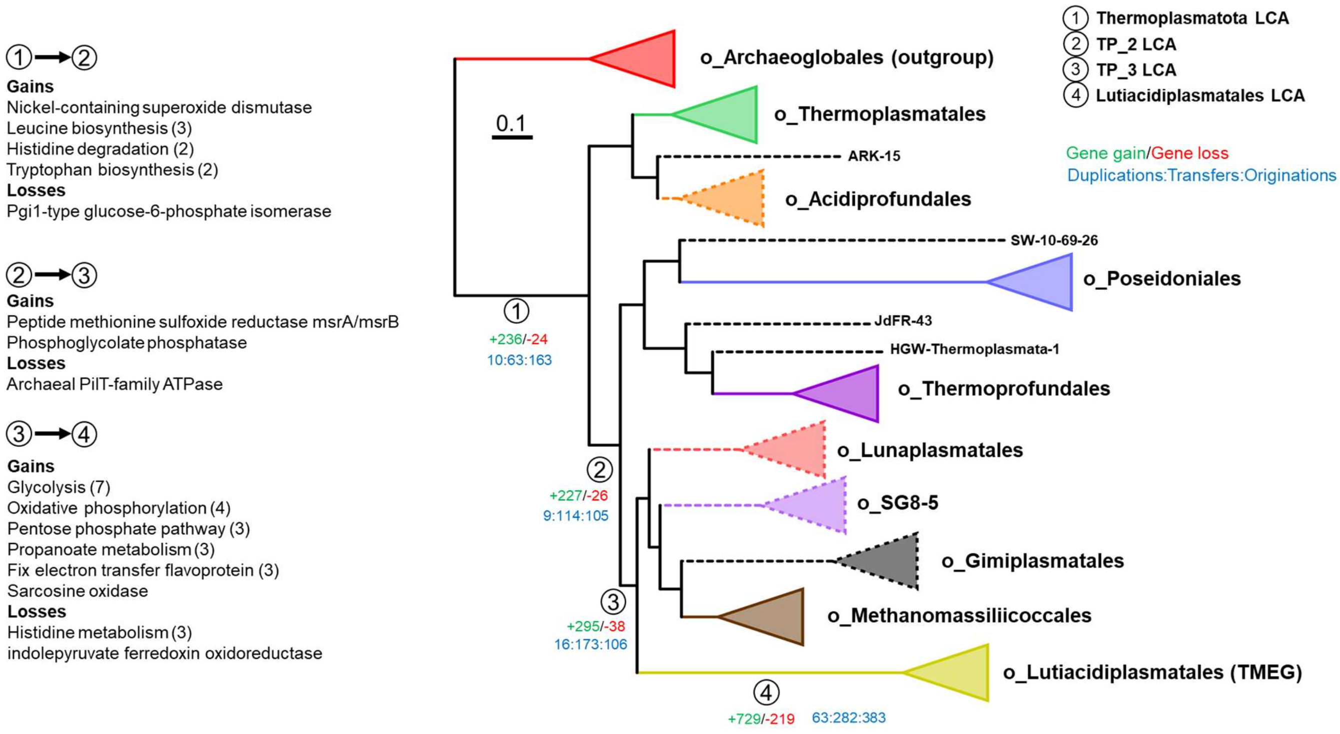


**Figure S26. Gene content evolution from the Thermoplasmatota LCA to Lutiacidiplasmatales LCA.** The gain and loss of gene families between the ancestral gene content reconstructions of the Thermoplasmatota LCA (1), the first divergence (2. TP_2 LCA), the second divergence (3. TP_3 LCA) and the final divergence to the Lutiacidiplasmatales LCA (4). The triangles represent collapsed clades, and the dotted triangles indicate that the clade consists of less than four representative genomes. The number of genes gained or lost for the listed metabolisms is conveyed in parenthesis.

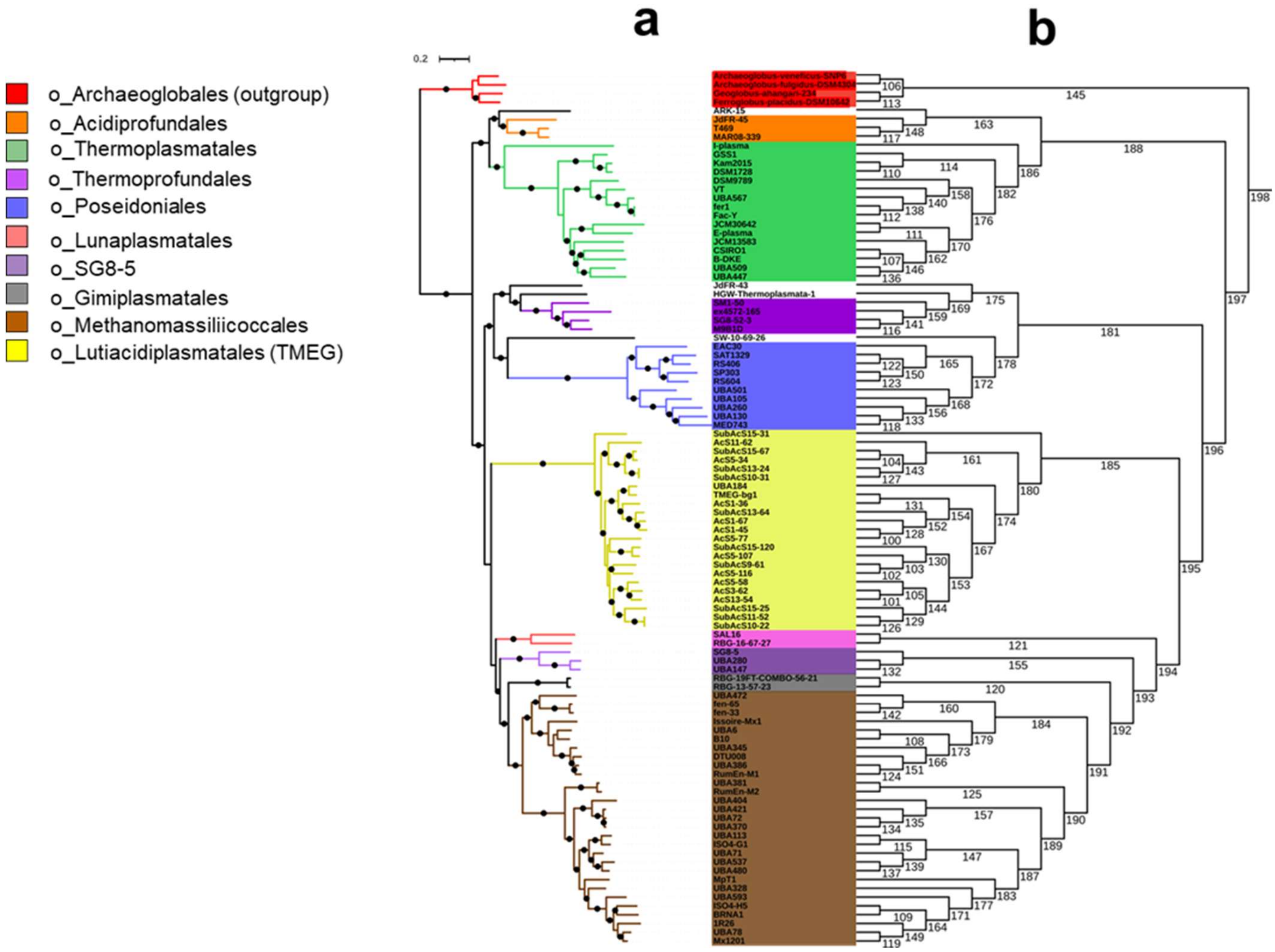


**Figure S27. Selected taxa (a) and branch labelled (b) phylogeny of Thermoplasmatota.** For the selected taxa tree (**a**), the ML tree was estimated using the best fitting model for each of the 71 concatenated genes. This species tree was used as input for the gene tree-species tree reconciliation. Dots indicate branches with >70% UFBoot and SH-aLRT support. The cladogram (**b**) possesses the topology of the species tree presented and is annotated with the branch numbers referred to in Supplementary Data 13. These two datasets are combined in Figure 3.
